# Alternative splicing fine-tunes modular protein interfaces to control protein localisation

**DOI:** 10.1101/2025.09.30.679652

**Authors:** Peter Kjer-Hansen, Karina Pazaky, Daisy Kavanagh, Helen E. King, Savannah O’Connell, Gabriela Santos-Rodriguez, Abigail K. Grootveld, Helaine Graziele Santos Vieira, Scott Berry, Javier Fernandez-Chamorro, Robert J. Weatheritt

## Abstract

Alternative splicing provides a pervasive means to expand proteome complexity, yet how it reorganises protein interactions and constrains where proteins act within cell remains unclear. By constructing an interface-resolved interaction network composed of 17,660 experimentally defined contact sites, we reveal that tissue-specific alternative splicing remodels protein connectivity by reshaping modular protein architecture. Longer exons reshape local interaction patterns whereas microexons fine-tune key interfaces linking distinct cellular processes. Integration of subcellular localisation data further indicates that such rewiring can redistribute proteins within cells. To test this, we developed a high-content imaging approach to systematically evaluate the influence of individual exons on protein localisation and screened a targeted library of protein isoforms differing in individual exons. 38% of the tested isoform pairs altered localisation, with microexons, although typically shorter than five amino acids, accounting for a substantial proportion of these effects. Bioinformatic and structural analyses identified that microexons can extend secondary structural regions and reposition charged residues, suggesting a potential to modulate local electrostatic environments. Consistent with this, biochemical analysis of a four–amino acid microexon in sorting nexin 2 - identified through our screen - confirmed that residue insertion, rather than side chain chemistry, was driving differences in protein localisation through repositioning of a flanking, charged residue. Together, these findings describe a principle by which alternative splicing fine-tunes interface architecture to coordinate protein assembly, localisation, and proteome organisation.

## INTRODUCTION

Almost all human protein coding genes are multi-exonic and their pre-mRNA therefore must be spliced to form mature mRNAs that can be translated^1^. The RNA splicing process is flexible, however, and variations in which parts of the pre-mRNA that are included and excluded from the mature mRNA, called alternative splicing (AS), occurs for nearly all spliced mRNAs^2,3^. Since AS can change the coding region of mRNAs, it can change the encoded proteins, although the contribution of AS to proteome diversity has been debated^4,5^. Recent efforts, however, have shown most well-expressed alternative transcripts give rise to protein isoforms detectable by mass spectrometry^6,7^, indicating the existence of vast AS-enabled protein diversity. While case studies have shown AS-derived proteins can impact practically every cellular process and are important for many biological processes^8–10^, the function of most isoforms remain unexplored. This indicates there are substantial amounts of the proteome yet to be explored, and increasing the understanding of the AS-derived protein isoforms holds both fundamental and applied value, including to therapeutics^11^.

High-throughput examination of AS-derived protein isoforms has mainly focused on protein-protein interactions (PPIs)^12–14^, especially binary PPIs^12,13^, which have shown AS frequently rewires protein interactomes, with most isoforms having distinct interactomes^12^. These analyses corroborate earlier bioinformatical analyses of tissue-regulated AS’s impact on proteins, which demonstrated AS is enriched in intrinsically disordered regions^15^ containing short linear interaction motifs^13,16,17^, which can mediate PPIs^18^. This suggests a common feature of alternative splicing is to enable a protein to function in new contexts via altered interactions, while maintaining its core functions^8^.

In multi-modular proteins, discrete binding motifs and globular domains combine in specific configurations that dictate molecular specificity, partner recruitment, and subcellular targeting^19–22^. Through the selective inclusion or exclusion of interaction modules, AS can potentially remodel these configurations and thereby modulate the assembly and dynamics of protein complexes. However, a systematic understanding of how such interface-level remodelling manifests across the proteome, and whether it contributes to the spatial and functional organisation of proteins within the cell, has yet to be established.

To address this, we combined interface-resolved structural network modelling with high-content imaging of protein isoforms, machine-learning-based analysis, and detailed biochemical and structural bioinformatics analyses to examine how alternative splicing fine-tunes protein connectivity, localisation, and molecular interfaces. Through this integrative framework, we reveal that tissue-specific splicing events preferentially fine-tunes the modular architecture of multi-domain proteins to enable local rewiring of interaction networks. These features are exon type-dependent with longer exons predominately affecting local interaction patterns, whereas microexons fine-tune key interfaces linking distinct cellular processes. Notably, we show that both exon types exert tangible effects at the cellular level with nearly half of the tested isoform pairs (16/42) displaying altered subcellular localisation. Structural and biochemical analyses further show that microexons within ordered regions, especially α-helices, influence protein architecture in two principal ways: by disrupting helical continuity to generate or extend loops, or by lengthening existing helices. Moreover, microexons within structured segments often occur near charged residues, and one way they can remodel interaction interfaces is by displacing these residues, as exemplified by two SNX2 isoforms identified in our imaging screen. Together, these analyses establish how alternative splicing fine-tunes the assembly and deployment of proteins across structural, spatial and networks levels.

## RESULTS

### Tissue-specific AS events refine interfaces within pathways whereas tissue-wide events regulate long-range interactions

To investigate how alternative splicing (AS) reshapes protein interactions at structural resolution, we constructed an interface-resolved human interactome in which each node represents a protein and each edge corresponds to an experimentally validated interaction. This network comprised 17,660 experimentally defined interfaces (**Table S1**), providing a detailed framework for assessing how exon choice influences protein connectivity (**Figure 1A**). Integrating differential splicing analysis^23,24^ across 75 tissues and cell types, we identified frame-preserving AS events that directly overlap mapped interaction sites, ensuring that only exons known to alter specific contact surfaces, rather than entire proteins, were considered. Approximately 6.5% of all interfaces were subject to splicing regulation (**Figure 1B**, **Figure S1A,** dPSI > 20, see Methods) defining a proteome-wide network of interface-level variation. Consistent with previous observations^13,16,17^, AS-regulated interfaces were enriched at high-degree modular hubs where interaction diversity is greatest (**Figure S1A**), indicating that splicing acts locally within highly connected neighbourhoods rather than globally rewiring the network.

**Figure 1.**
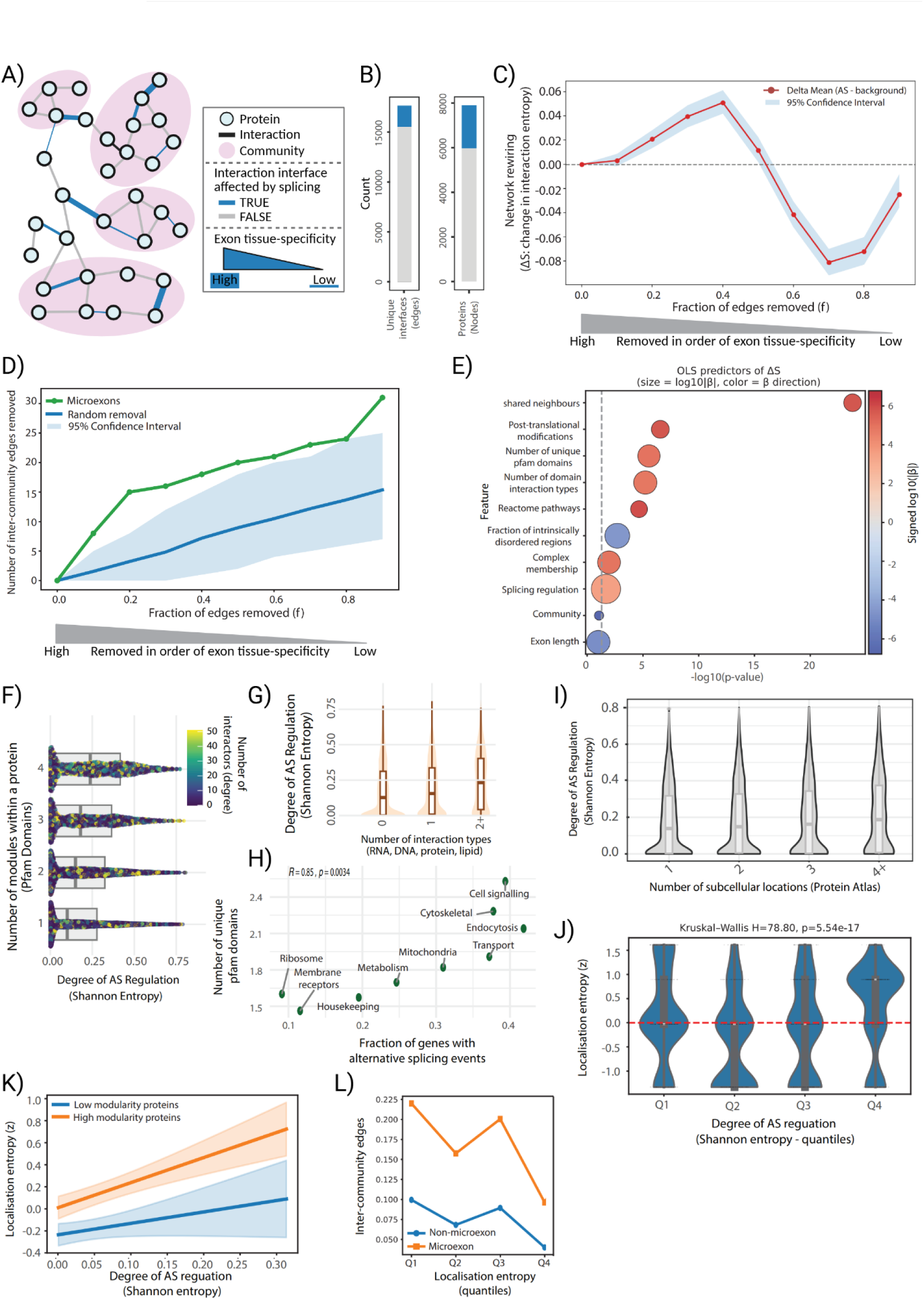
Alternative splicing can rewire protein networks and correlates with protein multi-modularity and diversification of protein subcellular localisation. **A)** Schematic of the interface-resolved protein interactome. **B)** Barplots showing the number of AS-regulated (blue) edges (representing interactions) and nodes (representing proteins) compared to non-AS-regulated (grey) edges and nodes within the interface-resolved interaction network. **C)** Percolation analysis of network entropy during progressive removal of AS-regulated edges. The red line indicates the mean difference in network entropy (Δ = SAS – Snull) as a function of the fraction of AS-regulated edges (f) progressively removed, from tissue-specific to tissue-wide events, with shaded regions showing the 95% confidence interval. The grey dashed line denotes Δ = 0 (no difference). Null background edges correspond to non-AS interactions randomly selected to match the degree distribution of AS-regulated edges. Positive Δ values indicate higher interaction diversity among AS-regulated interfaces. **D)** Percolation analysis of inter-community connectivity during progressive removal of microexon-regulated interfaces. The green line shows the number of inter-community edges removed as a function of the fraction of microexon-regulated edges (f) progressively removed from tissue-specific to tissue-wide events. Null edges correspond to non-AS interactions randomly selected to match the degree distribution of AS-regulated edge, which is represented by the blue line, with shaded areas indicating the 99 % confidence interval derived from 1000 permutations. Values above the null indicate that microexon-regulated interfaces disproportionately connect distinct functional communities. **E)** Regression analysis identifying protein features predictive of network entropy (ΔS). Bubble plot showing ordinary least-squares (OLS) model coefficients for top 10 structural, biophysical, and network features tested as predictors of ΔS. Bubble size reflects statistical significance (–log_10_ p-value) and colour indicates coefficient direction (blue = positive, red = negative). **F)** Violin and boxplots showing the distribution of splicing regulation (Shannon entropy) as a function of the number of Pfam domains per protein. Each data points represents a single protein, coloured by its total number of interaction partners (degree). **G)** Violin and Boxplots showing the distribution of splicing regulation (Shannon entropy) as a function of the number of interaction substrate types per protein (derived from domain annotation, see Methods). **H)** Scatter plot showing the mean number of unique domains compared to fraction of genes per functional group with detected alternative splicing events. **I)** Violin and boxplots showing the distribution of AS regulation (Shannon entropy) as a function of the number of annotated subcellular locations per protein (Human Protein Atlas). **J)** Violin and boxplots showing the distribution of localisation entropy as a function of the quartiles of AS regulation (Shannon entropy) **K)** XY plot showing the relationship between splicing regulation (x-axis) and localisation entropy (y-axis), with regression slopes plotted for proteins one standard deviation below (−1 SD) and above (+1 SD) the mean z-transformed number of Pfam domains (PFAMz), representing proteins with fewer and more modular domains, respectively. Shaded areas indicate 95% confidence intervals. **L)** A marginal effects plot showing the interaction between subcellular localisation diversity and microexon-containing proteins on inter-community connectivity. The y-axis indicates the model-predicted probability of an interface connecting distinct functional communities, plotted across increasing levels of subcellular localisation entropy (x-axis). Lines represent microexons-containing proteins (orange) and non-microexon-containing proteins (blue).

Because our goal was to understand how transcript diversity translates into cellular and structural rewiring, we next investigated whether alternative splicing reshapes interaction topology at the systems level. We therefore applied percolation analysis, in which AS-regulated edges are progressively removed to measure network robustness, and inter-community analysis to determine whether those edges bridge or occur within functional network communities. In this context, functional communities represent clusters of densely interacting proteins that often correspond to functional entities such as protein complexes, signalling cascades, or pathways. AS edges were removed in order of degree of regulation starting with tissue-specific events and ending with tissue-wide events (see Methods). This analysis revealed a sigmoidal change in the interaction diversity across the network, measured as network entropy (ΔS) (**Figure 1C**). Highly tissue-specific exons rewired interactions within their specific functional communities (**Figure 1C**, ΔS: p < 0.001) without affecting the connectivity across the whole network (**Figure S1B**, ΔE: p > 0.05). In contrast, removal of interactions regulated by tissue-wide AS events significantly affected interactions connecting discrete communities (**Figure S1C**, p < 0.002, empirical two-sided). These results suggests that while interfaces regulated by tissue-wide exons maintain long-range connections integrating different functional modules, tissue-specific exons rewire the network locally, acting as a local tuning mechanism rather than a disruptive force. This distinction indicates that tissue-specific splicing provides a means for context-dependent rewiring, which could potentially range from complex tuning to subcellular localisation changes.

Among tissue-specific events, microexons (< 30 nt) represent a striking class, as these events, in contrast to longer exons, frequently alter the structured domains of proteins^23^. We therefore next examined whether microexons displayed distinct patterns of regulation of the interaction network. In contrast to the previous observation of other tissue-specific exons, microexons were disproportionately regulating interfaces connecting distinct functional communities (**Figure 1D, Figure S1D-E**, p < 0.004, empirical two-sided) suggesting they regulated connections between pathways or complexes. Ordinary least-squares (OLS) modelling (Methods) further showed that inter-community connectivity depended on both tissue regulation and local topology, with a significant regulation × microexon interaction term (**Figure S1F**, p < 0.03, OLS t-test). This indicates that microexons represent a distinct subset of tissue-specific exons that preferentially occupy network positions capable of modulating crosstalk between different complexes or subcellular locations. Collectively, these observations suggest alternative splicing fine-tunes local interactions while preserving the broader integrity of the proteome.

### Multi-domain proteins distributed across multiple subcellular compartments are enriched for tissue-specific splicing events

To examine whether these network-level effects related to the biophysical properties of the proteins regulated by tissue-specific splicing, we next used regression and machine-learning models to identify protein features driving the network effects (Methods). This evaluation of over 20 separate features identified proteins with multi-domain structure^25^ that displayed interaction-substrate diversity (ability to engage protein, lipid, RNA, and DNA partners^25^) and distinct post-translational-modifications^26^ density (**Figure 1E**, all p < 10⁻⁵, OLS t-test), together with shared-neighbour connectivity, as the strongest predictors of ΔS. To determine whether these interface-level trends extend genome-wide, we analysed all frame-preserving, tissue-specific AS events irrespective of direct interface overlap. In line with the interface data, this analysis showed these events were disproportionately enriched in multi-domain proteins (**Figure 1F**, p < 3.47 x 10^-59^, Kruskal–Wallis test) that display interaction-substrate diversity (**Figure 1G**, p < 1.62 x 10^-16^, Kruskal–Wallis test). We next grouped proteins into broad functional categories chosen to represent a range of functional groups with differing splicing propensity, such as ribosomal, cytoskeletal, metabolic, signalling, and endocytic proteins. This comparison revealed a strong positive correlation between splicing propensity and median number of unique domains per a protein (**Figure 1H**, R = 0.85, p = 0.0034, Spearman’s rank). This suggests that AS acts preferentially on multi-domain hub proteins particularly associated with endocytosis and cytoskeletal organization that can accommodate local changes to their binding sites or domain arrangements potentially without disrupting overall protein stability or activity.

To explore whether these network-level effects correspond to spatial reorganisation within the cell, we integrated subcellular localisation annotations from the Human Protein Atlas^27^ and analysed all frame-preserving, tissue-specific AS events. Analysis of frame-preserving tissue-specific AS events identified a strong enrichment for these proteins to be localised across multiple cellular compartments (**Figure 1I**, p < 0.0029, Kruskal–Wallis test) suggesting that AS may broadly tune spatial deployment across the proteome. Indeed, proteins with tissue-specific events regulating interfaces exhibited higher localisation entropy (**Figure 1J**, all tissue-specific p < 5.543 x 10^-17^, Kruskal–Wallis across regulation bins) reflecting that tissue-regulated exons are preferentially expressed in proteins localised to multiple cellular compartments. This effect was most pronounced among highly modular proteins, where regulation strongly predicted localisation entropy (**Figure 1K**, β = 1.83, *p* = 9.5 × 10⁻¹³, OLS t-test), and remained significant when stratified by modularity (**Figure 1K, Figure S1G**, high-modularity: β = 0.58, *p* = 2.7 × 10⁻¹⁵, OLS t-test). Similarly, microexon-containing proteins displayed elevated localisation entropy (**Figure 1L**, β = 0.49, *p* < 0.001, OLS t-test), though this effect reversed when microexons occurred at inter-community interfaces (**Figure 1L**, β = –0.60, *p* < 0.05, empirical two-sided), suggesting microexon-containing proteins tend to have more specific subcellular niches. Together, these observations suggest that tissue-specific exons and microexons may remodel protein localisation through complementary routes, with tissue-specific events fine-tuning organisation within pathways, and microexons redirecting proteins at the junctions that connect distinct cellular processes.

### High-content imaging reveals prevalent localisation differences between isoforms

To determine whether these network-level effects translate into measurable changes in subcellular distribution, we next examined how alternative splicing influences protein localisation at the single-cell level using a high-content imaging approach. We focused on tissue-regulated AS events (both microexons and longer cassette exons) in multi-domain proteins involved in endocytosis and cytoskeletal organisation, which we found to display a high degree of tissue-specific AS regulation and high prevalence for multi-domain structures.

To do this we generated a protein isoform library for imaging analysis by sub-cloning open reading frames (ORFs) encoding distinct isoforms into plasmids fused with mCherry. For each gene, we cloned at least two isoforms that differed by a cassette exon. The final library comprised 68 isoforms across 32 genes, yielding 44 possible within-gene isoform comparisons (**Table S2**), of which only ∼16% had previously been examined for localisation differences^28–34^. Our aim was to determine whether isoforms from the same gene showed distinguishable subcellular localisation or behaviours (**Figure 2A**) based on quantitative features of the mCherry fluorescence signal, using identical and clearly distinct constructs as negative and positive controls.

**Figure 2.**
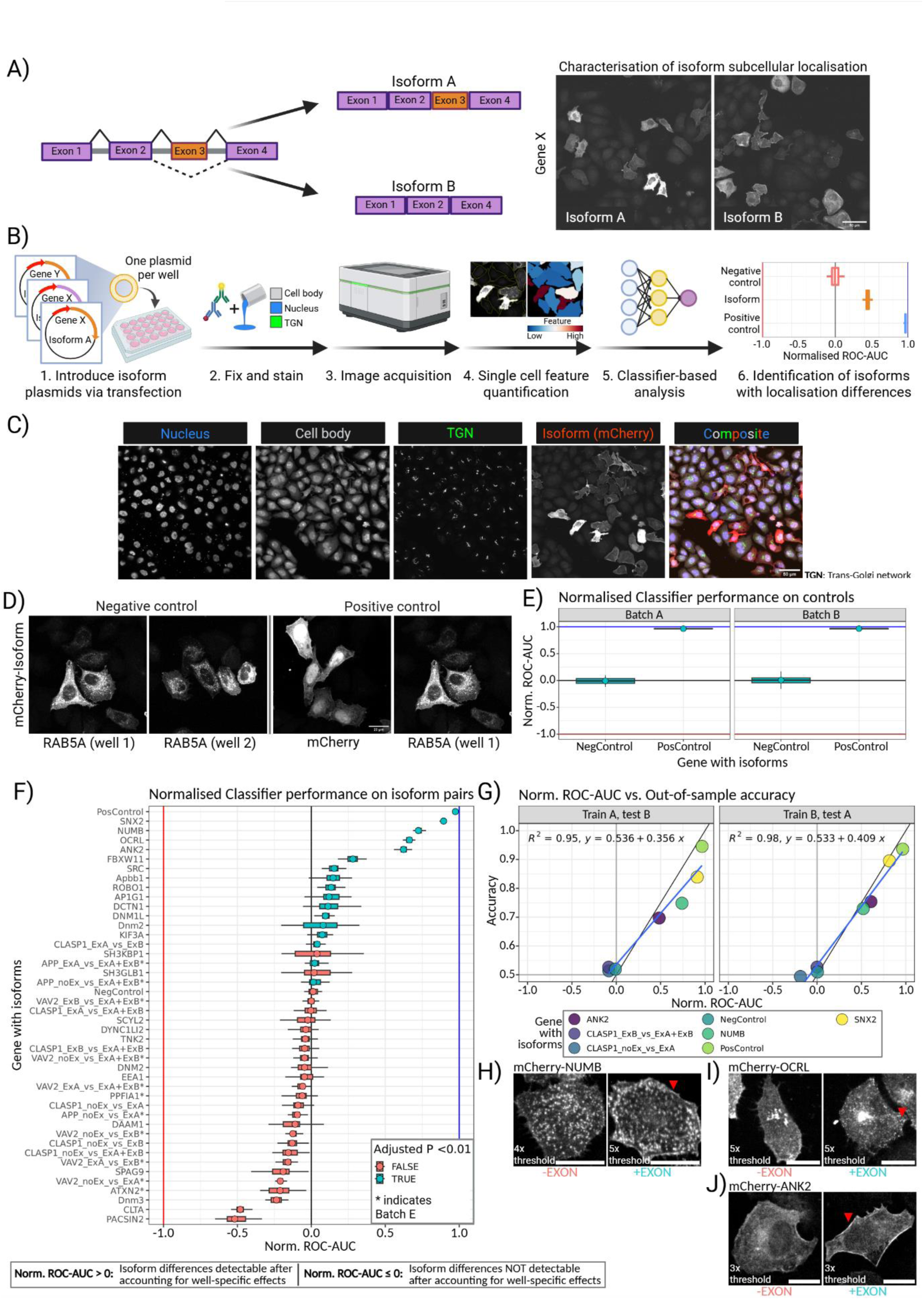
High-content imaging-based examination of a targeted protein isoform library reveals protein isoforms frequently differ in localisation. **A) Left**, schematic illustrating inclusion or exclusion of cassette exon in spliced mRNA leads to different transcript isoforms that can encode different protein isoforms. **Right**, example images of fluorescent protein isoforms expressed in HeLa cells. **B)** Workflow of high-content imaging and classifier-based examination of protein isoforms (see Methods). TGN, Trans-Golgi network. **C)** Example of the four channels acquired for a field-of-view. The focus slice is shown after illumination correction. Scale bar: 50µm. **D)** Example images of cells expressing the mCherry constructs used as positive and negative controls. The negative control consists of two separate wells transfected with the same mCherry construct (mCherry-RAB5A). The positive control consists of two separate wells transfected with mCherry and mCherry-RAB5A, respectively. Scale bar: 20µm. **E)** Boxplot showing the ability of classifiers trained on mCherry features measured in cells expressing control plasmids to correctly assign the plasmid ID (which corresponds to well ID), quantified as normalised ROC-AUC (Norm. ROC-AUC). Cells used for the examination came from three experimental repeats that were pooled for the classifier-based analysis with n cells ≥ 500 for each comparison after pooling. The experiment and analysis were performed twice (batch A, batch B), which were evaluated separately. The boxplots illustrate the normalised ROC-AUCs from each of 50 iterations of cell sampling and classifier evaluation, and dot indicates median normalised ROC-AUC (n = 50). Coloured lines indicate perfect (blue), neutral (black) or worst (red) possible classifier performance, respectively. **F) Top**, Boxplot showing the performance of classifiers trained on mCherry features of cells transfected with protein isoforms (or controls) making up isoform pairs, quantified as normalised ROC-AUC (Norm. ROC-AUC). All isoform pairs from batches A-E are shown. The cells used to examine a given isoform pair came from three experimental repeats from a given batch that were pooled for the classifier-based analysis. For isoform pairs present in multiple batches (batch A and B), cells expressing these were pooled prior to classifier-based analysis and so cells came from 6 experimental repeats. The shown controls also derive from pooling batch A and B. n cells ≥170 for each pair after pooling, with median n cells = 967, and numbers for each pair listed in **Supp. Table 3**. The boxplots illustrate the normalised ROC-AUCs from each of 50 iterations of cell sampling and classifier evaluation, and dot indicates median (n = 50). The isoform pairs are coloured according to whether the lower bound of a 99% bootstrap confidence interval included 0, after multiple testing correction (false discovery rate), with p-values < 0.01 being considered significant and indicating isoform differences. Batch E pairs are marked with *. Gene names in all caps indicate isoforms come from human, whereas genes only starting with upper case come from mouse. **Bottom,** interpretation of normalised ROC-AUC score. **G)** XY plot showing the correlation between the normalised ROC-AUC and the out-of-sample accuracy for the controls and five isoform pairs occurring in both batch A and B (n = 7), with pair identity indicated by dot colour. The normalised ROC-AUC for batch A is compared to the out-of-sample accuracy when trained on batch A and tested on B (vice versa for ROC-AUC for batch B). A linear regression was fitted to the data. After pooling cells across experimental repeats, ≥232 cells were present in both training and test set for a given comparison. **H)** Representative images of HeLa cells expressing mCherry-NUMB isoforms. The expression level (relative to the transfection threshold) is indicated in white text. Red arrowheads indicate the features that are accentuated in the +EXON isoform. Scale bar: 15µm. **I)** Representative images of HeLa cells expressing mCherry-OCRL isoforms. The expression level (relative to the transfection threshold) is indicated in white text. Red arrowheads indicate the features that are accentuated in the +EXON isoform. Scale bar: 15µm. **J)** Representative images of HeLa cells expressing mCherry-ANK2 isoforms. The expression level (relative to the transfection threshold) is indicated in white text. Red arrowheads indicate the features that are accentuated in the +EXON isoform. Scale bar: 15µm.

Isoform-encoding plasmids were transiently transfected into HeLa cells (**Figure 2B**), which were then fixed, stained, and imaged by automated confocal microscopy across four channels (DAPI, NHS-ester, anti-GOLGA1, and mCherry) (**Figure 2C**). Single cells were segmented, and ∼200 image-derived features describing the mCherry signal were extracted to identify transfected cells and quantify their fluorescence properties (**Figure S2**). For each isoform pair, measurements of mCherry signal features were compared using a classifier-based approach to identify differentially localising isoforms (**Figure 2B**). Briefly, for the classifier-based analysis, cells expressing each isoform within an isoform pair were pooled, down-sampled to equal numbers, and used to train a classifier with 10-fold cross-validation to assign isoform identity (**Figure S3**, Methods). Classifier performance was measured as mean ROC-AUC (0.5 = random; 1.0 = perfect separation) which we used to quantify the degree of difference between isoforms. Internal normalisation classifiers were trained on non-transfected cells from the same wells as the corresponding isoform pair to account for well-specific batch effects. The classifiers also assigned feature importance scores, which identifies the most discriminating signal characteristics between the isoforms.

To validate the approach, we assessed its performance on negative controls (duplicate mCherry-RAB5A wells) and positive controls (mCherry-RAB5A vs mCherry) (**Figure 2D**). As expected, classifiers trained on transfected cells matched their internal normalisation classifiers for the negative controls but showed near-perfect performance for positive controls (ROC-AUC ∼0.98 in two independent experimental batches, each batch consisting of 3 replicates) (**Figure S4A-B**). To correct for well-specific effects, we used the internal normalisation classifiers to define a normalised ROC-AUC metric, symmetrical around zero, where values >0 indicated isoform differences were detectable after accounting for well-specific effects (**Figure S4C**). Positive and negative controls scored near 1 and 0, respectively, across batches (**Figure 2E**). We then confirmed the robustness of the approach via cross-batch testing (train on one batch, test on the other): out-of-sample classifiers achieved ∼0.92–0.94 accuracy for positives and ∼0.5 for negatives (**Figure 4D**), and scores correlated strongly (r ≈ 1.0) with within-batch normalised ROC-AUC (**Figure S4E**). Together, these results validated the classifier-based framework and its normalised ROC-AUC metric as a sensitive and reliable measure of construct-dependent localisation differences.

We next applied the high-content imaging and classifier-based framework to our isoform library. Isoforms were processed in five transfection, staining, and imaging batches (A–E) (3 replicates per batch), with overlapping isoforms between A and B being pooled to increase cell numbers without negative affecting performance loss, as assessed by pooling the controls (**Figure S5A-B**). Each batch included positive and negative controls to assess consistency, which had expected normalised ROC-AUCs near 1 and 0, respectively, except for batch E, where a brighter NHS-ester staining led to an anomalously low score for the negative control (∼ –0.3) (**Figure S6A-B**). Despite this, we considered all batches suitable for analysis as E’s positive control performed as expected. In total, we successfully analysed ∼94% (64/68) of isoforms across 30 genes, yielding 42 pairwise comparisons (**Figure 2F, Table S3**). Normalised ROC-AUC values ranged from –0.5 to ∼1.0, with most between –0.2 and 0.2, indicating both distinct and similar localisations among isoform pairs. Significant differences (bootstrap 99% CI > 0) were detected in 38% (16/42) of pairs and the positive but not the negative control (**Figure S7**), with similar rates for microexons (∼38%, 10/26) and longer exons (∼38%, 6/16). Importantly, the normalised performance was independent of overall expression level differences between isoforms within pairs (**Figure S8A**, correlation test p = 0.76), and the fraction of total cells that were contributed by a single replicate (**Figure S8B**, correlation test p = 0.10). Moreover, classifier performances were highly consistent when we repeated the classifier-based analysis after separating cells within isoform pairs into low and high expression groups (**Figure S9A-B**), suggesting the isoform differences are present across all detectable expression levels, with the low intensity groups being only slightly above the mCherry detection limit (transfection threshold) for many isoform pairs (**Figure S9C**). These analyses all support the detected differences have biological rather than technical origins.

To confirm this, we performed out-of-sample testing on isoforms recurring in batches A and B. Classifier accuracy correlated strongly with normalised ROC-AUC (p = 0.01 and 0.003; slopes ∼0.36–0.41) (**Figure 2G**), consistent with true construct-based differences rather than well-specific effects. Notably, 71% (5/7) of isoform pairs with previously reported localisation differences^28–34^ were among those identified here (**Table S3**), providing external validation. Together, our imaging-based examination of isoforms therefore support evidence from our *in silico* analysis that alternative splicing frequently alters subcellular localisation of multi-domain proteins and does so via both microexons and longer exons in the examined isoforms.

### Orthogonal validation of localisation screen through literature and interaction analyses

To independently validate the accuracy of our isoform-specific localisation screen, we next closely compared our observed isoform localisation with the existing literature and then used orthogonal evidence to examine whether the observed subcellular localisation differences was associated with other changes, substantiating they reflect biologically meaningful regulation.

Our classifier-based analysis indicated marked differences between the NUMB and OCRL isoforms, and both have previously been examined in-detail^29,30^, which enabled us to assess whether the features that were important for classification and captured isoform differences (referred to as Isoform Distinguishing features, see Methods) matched the previously reported differences. The NUMB isoforms were distinguished by an Intensity_Edge feature (**Figure S10A-B**) - which describes the relative signal amount from the cell border - and we observed enrichment of the NUMB+EXON isoform at the plasma membrane (**Figure 2H**), consistent with previous reports^30^, while the OCRL isoforms differed in granularity and intensity variance features (**Figure S10C-E**), reflecting established vesicular versus diffuse distributions^29^, similarly reflected in our images (**Figure 2I**). These examples supported that the classifier-based approach identified features describing actual localisation differences rather than imaging artefacts, further supporting our results (**Table S4** contains Isoform Distinguishing features for all differentially localising isoform pairs). Notably, for both NUMB and OCRL, AS alters interfaces that modulate interactions with partners that likely explain their localisation patterns—membrane lipids for NUMB^30^ and clathrin for OCRL^29^ — exemplifying how interface modulation can underlie spatial redistribution.

Turning to orthogonal evidence, we reasoned that if AS-driven changes in localisation are functionally relevant, they should also manifest as differences in protein–protein interactions, since altered subcellular context constrains accessible interaction partners. Integrating data from LUMIER-based PPI screens^13,23,35^ we observed that isoform pairs with greater localisation divergence also exhibited greater interaction divergence (R² = 0.58, p = 0.027; **Figure S10I**, **Table S5**). This cross-assay concordance corroborates that isoform-dependent localisation changes detected by the screen correlates with underlying shifts in protein interaction networks, supporting the overall validity of the approach. A further complementarity in integrating interaction and localisation data is exemplified by the ankyrin-2 (ANK2) isoforms, whose localisation were previously uncharacterised. Via our imaging, we identified multiple isoform-distinguishing features, with intensity variance and radial distribution being the top features (**Figure S10F-H**) - indicative of altered enrichment at the cell periphery – and we confirmed the ANK2+EX isoform was enriched at the membrane (**Figure 2J**). The examined exon in ANK2 overlaps a ZU5 domain, and the previous interaction study showed that ANK2+EX interacts more strongly with cytoskeletal and membrane-associated proteins than ANK2–EX^13^. This suggests a close correspondence between AS-modulated interfaces, altered interaction potential, and subcellular localisation in ANK2 and, underscores the complementarity of these orthogonal approaches in isoform characterisation, which aligns with recent findings for transcription factor isoforms^36^.

### Microexon inclusion displaces charged residues within secondary structure elements

Our imaging-based screen revealed that microexons, despite encoding only a few residues, alter protein localisation to a similar extent as longer cassette exons. This is notable because alternative splicing typically modulates localisation and interaction networks through well-characterised mechanisms such as domain inclusion^37,38^ or the gain or loss of short linear motifs^16,39^, which adds or removes interaction interfaces. By contrast, microexons are very short insertions that often occur in structured regions and adjacent to, or partially overlapping, existing interfaces^23^. This raises the question of how such small insertions, which are unlikely to destroy or generate entire interaction surfaces, can still exert consistent functional effects. One possibility is that microexons act through subtle structural mechanisms by adjusting local geometry or the positioning of key residues within preexisting folds and interfaces. To explore this, we examined whether microexons display properties that allow them to preserve or modulate protein secondary structure, which may be one way they can alter the architecture and interfaces of structured regions.

Since the top microexon hits in our imaging screen (SNX2, OCRL) were located within or adjacent to α-helices, we first examined whether microexons in general show a bias towards preserving helical structural architecture. We analysed the nucleotide length distribution of microexons in coding regions, comparing microexons in general to those predicted to occur within α-helices (**Figure 3A**). We observed microexons of 12nt and 21nt, encoding 4 and 7 residues, respectively, corresponding to one and two helical turns, were enriched in predicted helices (**Figure 3B**). This suggests that short microexons often preserve helical structure, although enrichment of slightly longer 24nt and 27nt microexons implies this bias may be limited to shorter insertions. We next examined the position of microexons within predicted helices (**Figure 3C**). Interestingly, helix-preserving microexons (12nt, 21nt) were rather evenly distributed along helices, whereas those with potentially disruptive lengths were enriched near helix ends, where they may extend or induce loops. This pattern was exemplified by AlphaFold3^40^ models of ERC1 and TEAD1, where microexon inclusion is predicted to either maintain helical geometry or subtly change it by forming/extending loops, respectively (**Figure 3D**). To explore the biochemical consequences of these occurrences, we compared the amino acid compositions of microexons and their flanking residues, with those of longer (>30nt) tissue-specific cassette exons with similar regulation patterns. Microexons were enriched in helix-promoting residues and charged residues, regardless of whether they occurred in predicted α-helices or disordered regions (**Figure 3E**), although this trend weakened when β-strands and loops were considered. Moreover, residues flanking microexons within structured regions contained more charged residues than those surrounding longer exons, a difference not observed for disordered regions (**Figure 3E**).

**Figure 3.**
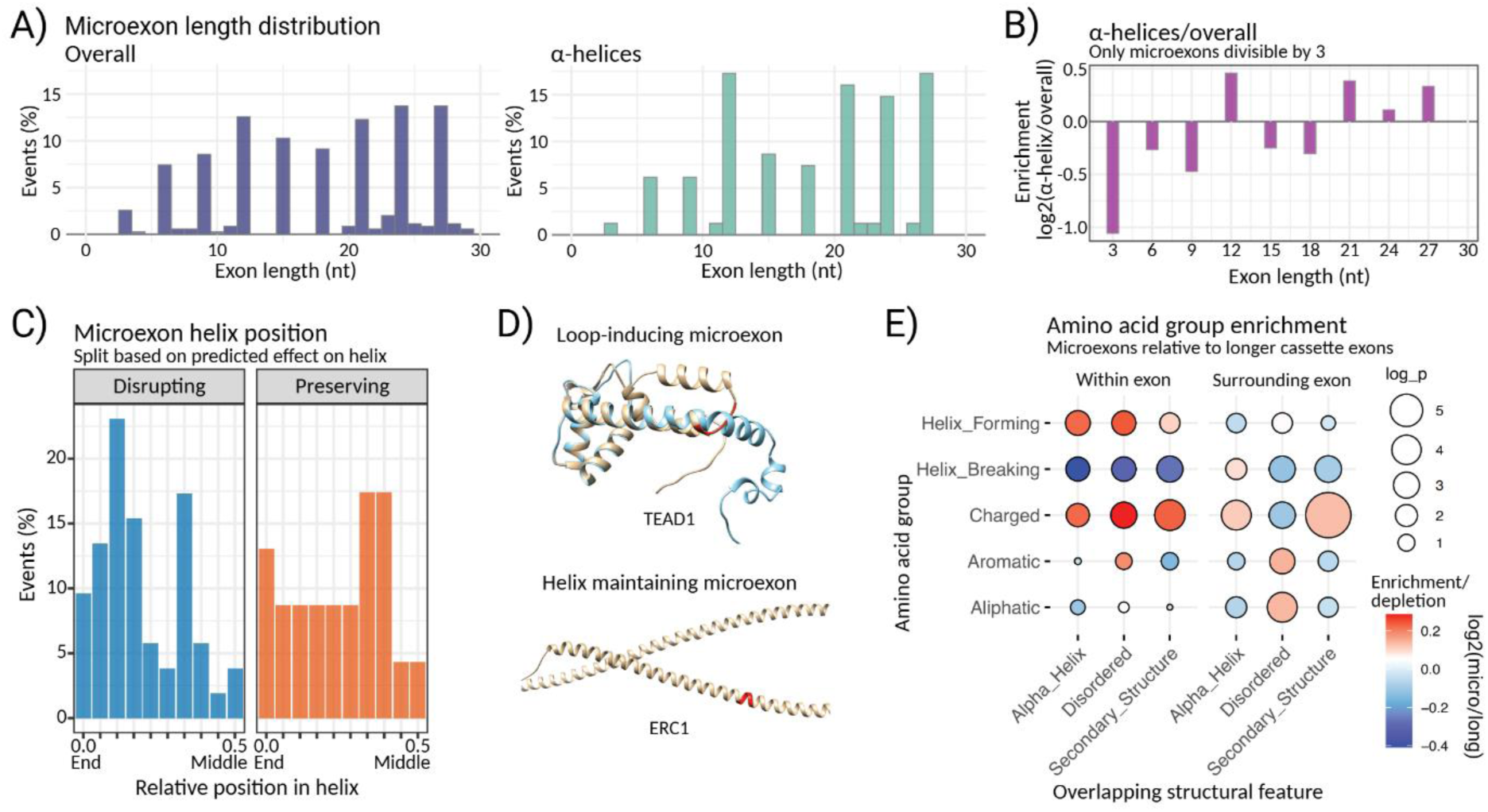
Microexons potentially affect α-helical structures in two common ways. **A)** Relative frequency histogram of microexon nucleotide (nt) lengths for tissue-specific microexons (<30nt) (Methods), **Left**, overall. **Right**, microexons predicted to overlap α-helices. n microexons = 353, n microexons in α-helices = 82. **B)** Barplot showing the enrichment and depletion of alternatively spliced microexons with specific lengths in α-helices, calculated as 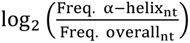. Only microexon lengths divisible by 3 are included. **C)** Relative frequency histogram of tissue-specific microexon positions within predicted α-helices, quantified as the relative distance of the point where the microexon is inserted to the nearest α-helix end (Methods). Microexons are grouped according to whether their length is predicted to preserve (12nt, 21nt) (Preserving) or disrupt (other lengths) the helical structure (Disrupting). **D)** AlphaFold structures of TEAD1 and ERC1 isoforms, showcasing microexons predicted to induce/extend loops in alpha helical structures and microexons predicted to preserve the helix. Microexons are coloured red. For TEAD1, isoforms with and without the microexon are shown in separate colours (blue and gold) and overlaid. **E)** Bubble plot showing amino acid enrichment within exons (exon) and flanking regions (surround), for tissue-specific microexons overlapping predicted α-helices, disordered regions, and secondary structure elements (α-helices, β-strands, loops), compared to longer cassette exons (≥30nt) with a similar inclusion pattern. Only exons divisible by three were included. Amino acids were grouped according to properties in a non-mutually exclusive manner: helix forming: L, I, F, E, Y, W, M, helix breaking: P, G , Charged: E, D, K, R , Aromatic: F, Y W, H, Aliphatic: L, I, V. Colours indicate degree of enrichment (calculated as 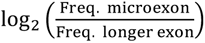, size indicates statistical significance calculated by fisher-exact test. Flanking residues were the first 30 residues upstream, and 30 residues downstream of the microexon. n microexons in α-helices = 77.

Together, these observations suggest microexons may affect basic α-helical architecture in at least two ways: by preserving and extending helices or by disrupting the helix to some extent, including forming/extending loops. When they occur in α-helices and other secondary structure (β-strands and loops), they may displace the flanking residues, notably charged residues, which potentially serves as a way whereby microexons can fine-tune the properties of preexisting interfaces and possibly modulate intermolecular interactions.

### Insertion of the SNX2 microexon, not its composition, drives localisation and interaction changes

To test the predictions from this bioinformatic analysis for α-helical microexons, we next examined the two isoforms of sorting nexin 2 (SNX2), which were the top hit from our subcellular localisation screen and differed in an α-helical microexon. SNX2 is a multi-domain protein that regulates retrograde transport and recycling of internalised membrane protein^41^, and the isoforms were previously uncharacterised. Our imaging data showed that microexon inclusion markedly altered SNX2 localisation with approximately 20 features being isoform-distinguishing (**Figure S11A**), the most important one relating to granularity (**Figure S11B**). The granularity feature captured the more punctuated, vesicular localisation of SNX2-EX which at higher expression formed aggregates – previously described as swollen endosomes for GFP-SNX2-EX^42^ - whereas SNX2+EX displayed a more diffuse cytoplasmic distribution at all expression levels (**Figure 4A-B**). The pronounced localisation differences between the isoforms highlighted the SNX2 isoforms as a model setting to examine how an α-helical microexon can tune domain geometry and residue positioning with consequences for localisation.

**Figure 4.**
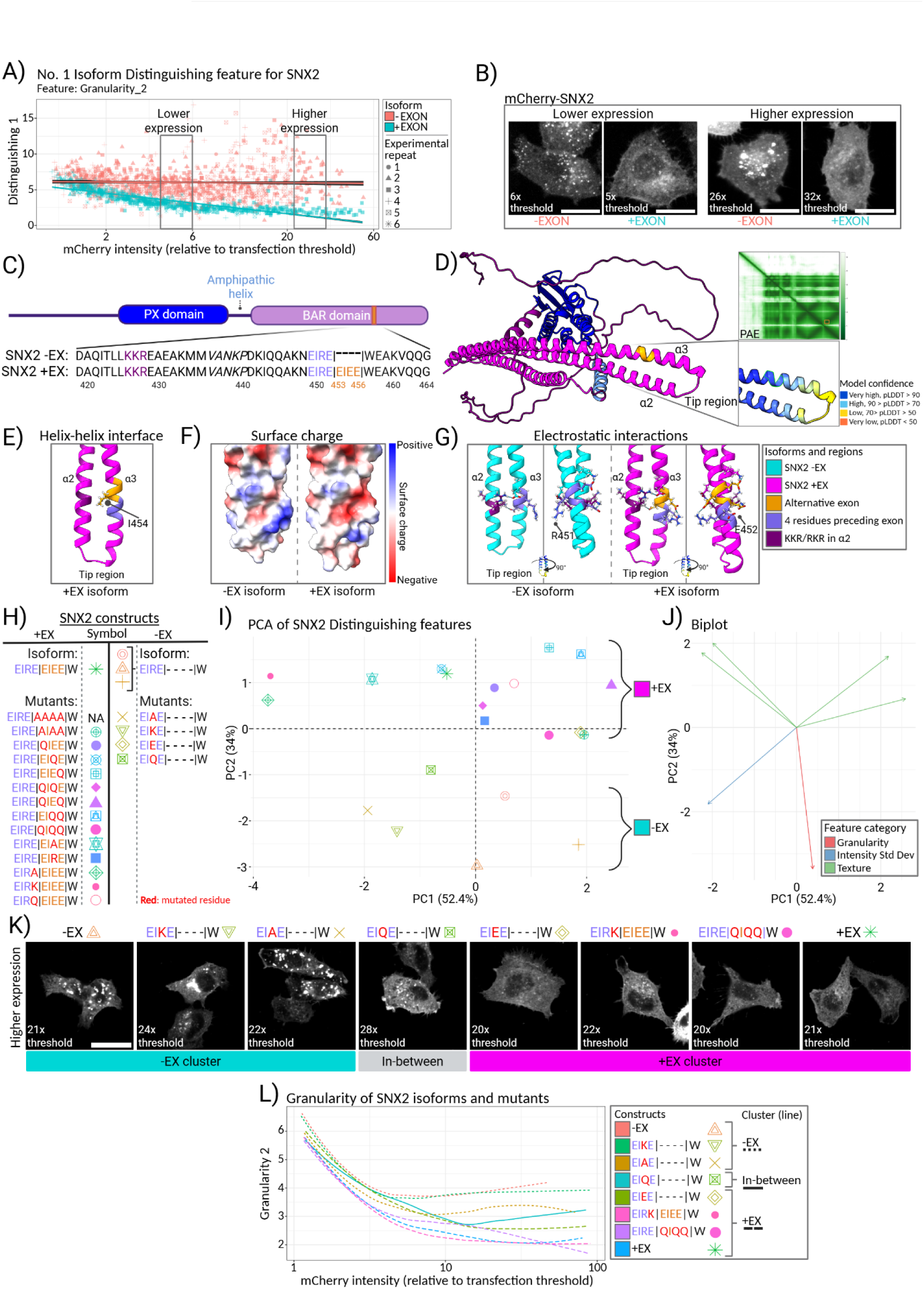
SNX2’s subcellular localisation is altered by insertion of the microexon, not by the composition of its residues. **A)** XY plot of Top 1 Isoform Distinguishing Feature for the SNX2 isoforms (mCherry Granularity) against cell mean mCherry intensity (relative to transfection threshold). Each data point corresponds to a cell with colour and shape indicating SNX2 isoform identity and experimental repeat ID, respectively. Cells come from N = 6 experimental repeats, with pooled n cells > 1300. Separate linear regression lines were fitted to SNX2+EX and SNX2-EX-expressing cells. Two rectangles (lower expression, higher expression) indicate where the representative cells (Figure 4B) fall within the overall population. **B)** Representative images of HeLa cells expressing mCherry-SNX2 isoforms. Cells with two different expression levels (lower, higher) are shown with the expression level (relative to the transfection threshold) indicated in white text. Scale bar: 15µm. **C)** Primary structure of human SNX2 with structured domains indicated as ovals and disordered regions as lines. The microexon (orange) is inserted into SNX2’s BAR domain (purple), and the exon’s residues (orange) as well as the residues surrounding the exon are indicated (residues 419-464). The four residues preceding the exon are lavender, and the KKR motif (426-428) is dark purple. Residue numbers prior to the exon corresponds to Uniprot ID: O60749. Residues after the exon have a position number that’s +4 that of Uniprot ID: O60749. **D)** AlphaFold2 (AF2) structure of human SNX2+EX. The BAR domain is coloured magenta, the amphipathic helix is light blue, and the PX domain is dark blue. The microexon is coloured orange and inserts as an additional helical turn in the α3 helix of the BAR domain. The α2 helix and tip region connecting α2 and α3 are indicated. The predicted aligned error (PAE) for the model (region with exon marked by orange square) and the predicted local distance difference test (pLDDT) for the BAR region with the microexon is shown (exon marked with green outline). **E)** AlphaFold2 model showing the position of I454’s sidechain, encoded by SNX2’s microexon, for the human SNX2+EX isoform in the BAR region containing residues 419-464. The BAR domain is viewed from the side that faces lipid membranes. The microexon is orange, the four residues preceding the exon are lavender, and the KKR motif (426-428) is dark purple. **F)** Surface charge of AF2 models of human SNX2 isoforms in the BAR region containing residues 419-464. The BAR domain is viewed from the side that faces lipid membranes. **G)** Predicted electrostatic interactions between residues in the AF2 models of human SNX2 isoforms in the BAR region containing residues 419-464. Residues engaging in electrostatic interactions (salt bridges) are shown as well as residues that form electrostatic interactions in one isoform but not the other (R451, E452). The microexon, the four preceding residues, and the KKR motif are indicated. The left panel shows the BAR domain from the side that faces lipid membranes. **H)** List of SNX2 mutant constructs examined via high-content imaging. Residue colours correspond to colours in Figure 4C. Mutated residues are indicated in red. Each construct has a symbol that indicates their position in the PCA. Note, three technical replicates for SNX2-EX were included. **I)** XY plot of PC1 versus PC2 for principal component analysis (PCA) of the mCherry-SNX2 construct imaging data using the population means of the SNX2 Isoform Distinguishing features (after highly correlated features were removed) using the transfected cells as input. Two clusters that are perpendicular to the granularity vector are indicated, and these contain the -EX and +EX isoforms, respectively. Cells were pooled across three experimental repeats (N=3), with ≥185 cells expressing each construct (cell numbers for each construct are shown in **Figure S13**. **J)** Biplot of the Isoform Distinguishing features, where arrow directions show how the features contribute to each PC axis (PC1, PC2), and arrow length indicates their importance. Arrow colours reflect feature classes with granularity shown in red. **K)** Representative images of HeLa cells expressing mCherry-SNX2 isoforms and select mutants. The expression level (relative to the transfection threshold) for the brightest cell in each image is indicated in white text. The PCA symbol and cluster identity for each construct is indicated. Scale bar: 25µm. **L)** Line plot of mCherry Granularity (Y-axis) against cell mean mCherry intensity (relative to the transfection threshold) for mCherry-SNX2 isoforms and select mutants that represent the different clusters and have representative images (Figure 4K). The lines are fitted to all transfected cells expressing the specified constructs, pooled across three experimental repeats (N=3), with ≥185 cells expressing each construct (cell numbers for each construct are shown in **Figure S13B**). Line colours reflect construct identity and line type indicates PCA cluster.

SNX2’s microexon encodes four residues (EIEE) that inserts into SNX2’s BAR domain (**Figure 4C**) - a coiled-coil structure that mediates lipid binding, dimerisation, and higher-order oligomerisation^43^. To understand how this altered SNX2 function, we first modelled both isoforms using AlphaFold2^44^. This suggested the microexon inserts into the BAR domain as a single additional helical turn in the α3 helix (**Figure 4D)** and presumably maintains the domain fold with the inserted I454 forming helix-helix interactions (**Figure 4E**), which are critical for coiled-coil stability^45^. The insertion made the surface charge more negative (**Figure 4F**) and was predicted to alter electrostatic interactions for a few, immediately flanking residues (R451/E452) (**Figure 4G**), through a combination of the exon’s three negatively charged glutamates and repositioning existing residues.

These observations were supported by sequence alignment across vertebrates of the residues in the microexon-containing region of SNX2’s BAR, which showed a high degree of conservation of both the microexon’s residues and the flanking residues (**Figure S12A**), and by AlphaFold2 models of both isoforms in select species (rat, chicken, zebrafish), which closely resembled the human isoforms (**Figure S1**2**B-D**). Collectively, this indicated that the microexon alters SNX2‘s BAR domain interface by changing surface charge, electrostatic interactions, and subtly repositioning of flanking charged residues, rather than changing the overall fold, and these effects may cause the localisation differences.

To test these structural predictions, we generated 18 mCherry-SNX2 mutants (**Figure 4H**), targeting the microexon’s residues and the flanking charged residues R451 and E452 with predicted electrostatic interaction changes, to explore the role of charge and electrostatic interactions in localisation. We retained I454 in all but one mutant because of its putative structural role. We transfected the mCherry-SNX2 constructs into HeLa cells and imaged them by automated confocal microscopy. All mutants, except the I454 mutant, expressed at comparable levels to the mCherry-SNX2 isoforms (**Figure S13A-B**) and we used these for subsequent localisation analysis. Here, we performed principal component analysis (PCA) of SNX2’s isoform-distinguishing features (after removing highly correlated features, Methods), which revealed two distinct clusters separating along the granularity feature vector (PC2) corresponding to SNX2−EX and SNX2+EX (**Figure 4I–J, Figure S13C**). This was consistent across replicates and absent in non-transfected controls (**Figure S13D–E**) supporting that the clustering reflected construct-specific rather than batch effects. By examining cells expressing constructs in either cluster, we confirmed the clustering indeed captured the isoform-specific granularity behaviour with aggregates forming in the -EX but not +EX cluster (**Figure 4K-L**). Interestingly, the clustering pattern reflected the presence or absence of the four-residue microexon with all +EX mutants clustering with the +EX isoform regardless of amino acid substitutions. This indicates that residue insertion, rather than the specific side chain properties, were driving the localisation differences. For -EX, the progressive substitution of R451 from positive or small to neutral to negative residues (R451K/A → R451Q → R451E) gradually shifted the constructs from the –EX to the +EX cluster, implicating the importance of positive charge at this location (451). Together, these results demonstrate SNX2’s microexon exemplifies an α-helical microexon that extends structure and that it alters SNX2 localisation by repositioning existing charged residues through residue insertion in a manner that is independent of its own amino-acid chemistry creating a new interface.

### Microexon-dependent residue displacement fine-tunes SNX2 localisation and interactions

To determine if the microexon’s effects extend beyond over-expression systems and localisation, we next examined whether the isoform-specific localisation differences occurred at endogenous levels and whether they were linked to changes in SNX2’s interaction network. To this end, we generated SNX2-knockout HeLa cells (**Figure S14A**) and then stably re-introduced untagged -EX or +EX SNX2 isoforms, as confirmed by western blotting (**Figure S14B**) and end-point RT-PCR (**Figure S14C**). We immunostained the cells with a pan-SNX2 antibody alongside HeLa wildtype cells (which express SNX2-EX (**Figure S14C**)), and imaged them using confocal microscopy, which confirmed the more granular, punctuated localisation of SNX2-EX compared to SNX2+EX at endogenous levels (**Figure 5A-B**) and over-expression (**Figure S14D-E**). Transfecting mCherry-SNX2 isoforms into the knockout cells reproduced these localisation differences (**SFig 14F**), demonstrating the mCherry-SNX2 behaviours capture SNX2 isoform behaviours at lower expression levels. Notably, at higher expression, we only observed aggregate formation for SNX2–EX fused to mCherry, suggesting that the observed aggregates are fusion-dependent, likely arising from the tag accentuating an underlying isoform-specific behaviour. Collectively, these results demonstrate SNX2’s microexon causes SNX2 to localise in a more diffuse, less vesicular manner, with the effect existing at endogenous expression levels in untagged SNX2.

**Figure 5.**
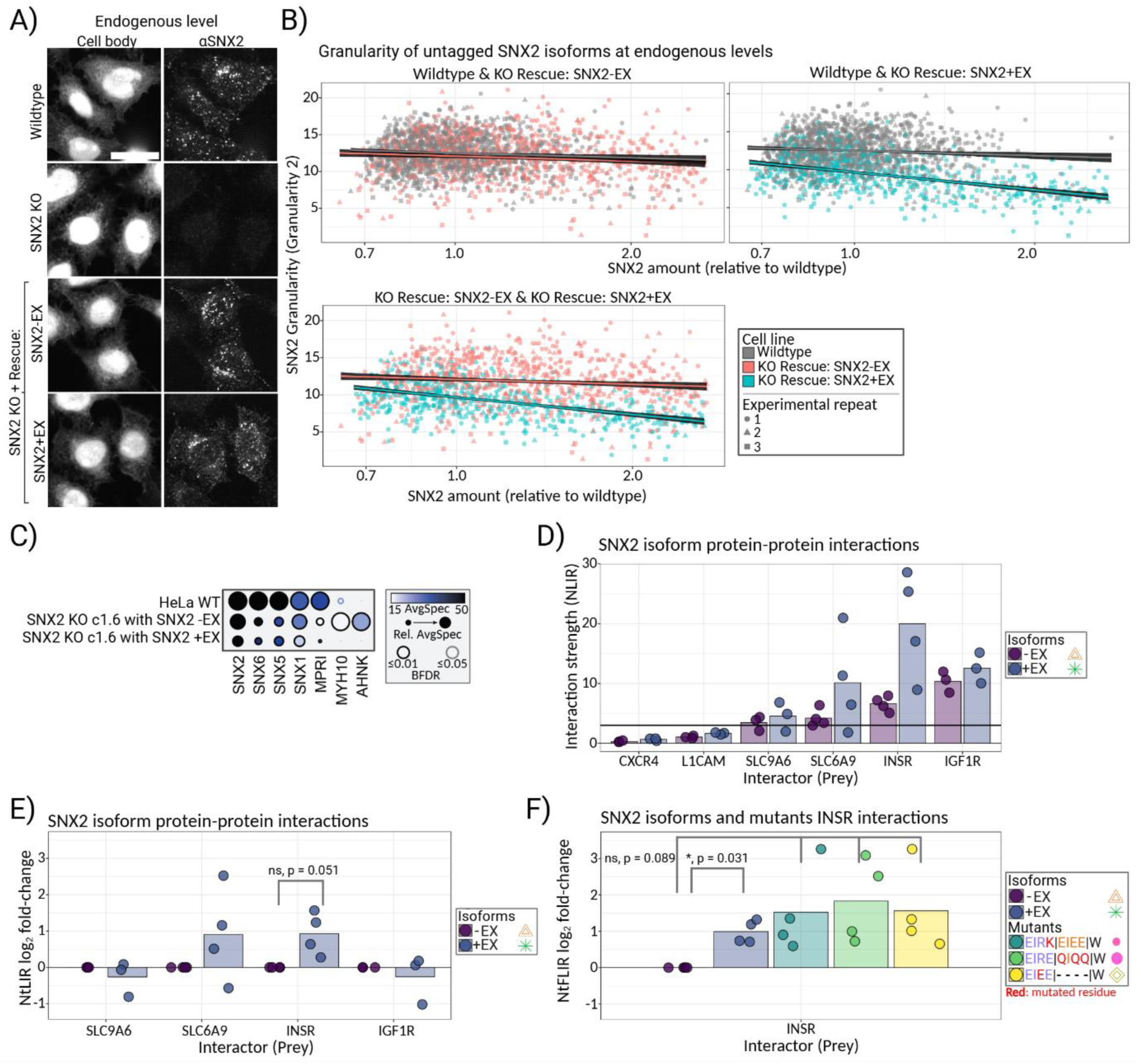
SNX2 isoforms localise differentially at endogenous levels and display different interactions with select protein partners, which relates to residue insertion rather than amino acid composition. **A)** Representative images of HeLa cell lines (wildtype, SNX2 KO, SNX2 KO + rescue stably expressing SNX2+EX or SNX2-EX) expressing SNX2 at endogenous levels with SNX2 detected via immunostaining. Scale bar: 25µm. **B)** XY plots of SNX2 amount (SNX2 intensity relative to median SNX2 intensity in HeLa wildtype) against SNX2 granularity 2 in the HeLa lines, at endogenous levels. Each point represents a single cell, colour represents cell line (n wildtype = 1351, Rescue SNX2-EX = 736, Rescue SNX2+EX = 558), and shape represents experimental repeat (N = 3). A linear regression line (x ∼ y) is fitted to the data, with the 95% confidence interval indicated as a black shade. Cell lines shown pairwise to reduce overplotting. **C)** Bubble plot of proteins detected upon affinity purification mass spectrometry (AP-MS) analysis of HeLa wild type and HeLa SNX2 isoform rescue lines. Only proteins with average spectral counts ≥15 are included. n = 3 independent experiments. Colours indicate average spectral counts, size indicates relative spectral counts, and line indicates Bayesian False Discovery Rate (BFDR). **D)** Barplot showing LUMIER interaction scores for SNX2 isoforms with 6 transmembrane proteins that are suggested to interact with and/or be regulated by SNX-BAR complexes. Interaction strength is quantified as NLIR (normalised LUMIER interaction ratio), and an NLIR>3 (black line) is considered a positive interaction^46^. Column height indicates mean NLIR across experimental repeats (n = 3 or 4 independent experiments), and dots reflect NLIR from each experiment. Colours reflect SNX2 isoform identity. **E)** Barplot showing LUMIER interaction strength of SNX2 +EX with the 4 transmembrane proteins that were identified as positive interactors (Figure 5D), relative to the SNX2-EX isoform’s interaction with them (log_2_ fold-change). This is the same data used for Figure 5D but, the interaction score is normalised to both the level of prey and bait protein (NtFLIR). Column height indicates mean log_2_FC across experiments (n = 3 or 4 independent experiments), dots reflect log_2_FC for individual experiments. Colours reflect SNX2 isoforms. Statistical test is one-sample t-test against µ = 0 (two-sided), p-value > 0.05, ns. **F)** Barplot showing LUMIER interaction strength of SNX2+EX and three SNX2 mutants with INSR, relative to the SNX2-EX isoform’s interaction with INSR (log_2_ fold-change). Individual dots reflect log_2_FC for individual experiments, and column indicates mean across experiments (n = 4 independent experiments). Interaction differences were tested using one-sample t-tests against µ = 0 (two-sided). p-values were corrected for multiple testing using false discovery rate. P-value < 0.05, *. P-value > 0.05, ns.

To assess whether these structural and localisation changes affected protein interactions, we next compared PPIs between SNX2 isoforms and select SNX2 mutants using affinity-purification mass spectrometry (AP-MS) and LUMIER assays. We performed AP-MS of SNX2 in the HeLa SNX2 rescue lines stably expressing either isoform (untagged), which showed both isoforms interacted with canonical SNX-BAR partners (SNX1, SNX5, SNX6)^47,48^, and the cargo MPRI/IGF2R^49^ (**Figure 5C**), confirming intact SNX-BAR complex formation. We then performed LUMIER to quantify interactions with SNX-BAR cargo proteins, examining interactions with six transmembrane cargos. We found both isoforms interacted with most cargos to a similar extent (**Figure 5D-E**), however, SNX2+EX had an approximately two-fold stronger interaction with the insulin receptor (INSR) (**Figure 5E-F**). Crucially, this enhanced binding persisted in two +EX mutants and the R451K -EX mutant (**Figure 5F)**, all of which clustered with SNX2-EX in the imaging experiment and did not form aggregates. The 4-residue insertion—not its amino-acid identity— was therefore sufficient to alter interaction strength. Collectively, these findings demonstrate that inclusion of the conserved SNX2 four-residue microexon is sufficient to reposition charged residues within the BAR domain, subtly altering electrostatics to fine-tune both subcellular localisation and protein-protein interactions, exemplifying one way microexons can alter structured regions.

## DISCUSSION

The extent to which alternative splicing regulates proteome organisation and function remains incompletely understood^8^, despite increasing evidence that protein isoforms are both prevalent and stably expressed^50–52^, and AS can have major implications for protein functions and interactions^12,36,53–56^. In this study, we reveal a common feature for alternative splicing is to act as a fine-tuning mechanism that by modulating proteins’ residue composition can rewire protein networks and subcellular localisation, for instance through microexons splicing which changes the electrostatic properties at individual interaction interfaces. Within tissues, such fine-tuning enables proteins to preserve their core molecular functions while acquiring context-specific properties that modulate specialised interactions, whereby isoforms can contribute to cellular identity and physiology^8,57,58^. An example of this is the gastric epithelium-specific cassette exon in MLCK1 which promotes tight junction localisation which enables this ubiquitously expressed protein kinase to regulate gastric barrier integrity through isoform-specific interactions^59^. Such tissue-specific behaviours can be disease relevant, with excessive tight junction localisation and activity of MLCK1 contributing to gastric barrier dys-function in inflammatory bowel disease^59,60^. Moreover, the barrier dys-function can be alleviated by preventing MLCK1’s tight junction localisation through small-molecules that blocks the isoform-specific interaction^60^. Fine-tuning of protein behaviours, including altered subcellular localisation, via tissue-specific AS can therefore be relevant to both health and disease and provide new therapeutic directions^11^, emphasizing the importance of studying these properties of AS.

Our network analysis revealed that AS preferentially targets multi-domain, multifunctional proteins containing binding surfaces for proteins, RNA, DNA or lipid partners. The multi-modular nature of multi-domain, multifunctional proteins likely makes them particularly amenable to regulation by AS, which can selectivity prune or enhance specific interfaces while preserving others^8,61,62^. By resolving these effects at the interface rather than gene level, our approach captures how AS locally modifies connectivity while maintaining global network integrity, supporting AS in fine-tuning protein function. Notably, this approach enables us to find that longer tissue-specific exons predominantly act within functional modules, likely acting as digital switches that add or remove individual interactions, whereas microexons operate at inter-community interfaces fine-tuning, but rarely removing, key connections. This dual mode of regulation provides a mechanistic basis for how alternative splicing can simultaneously refine local specificity and maintain global coordination across the proteome.

The combined observation that alternative splicing maintains network integrity while targeting multi-modular proteins distributed across multiple subcellular compartments prompted us to develop a high-content imaging approach to test whether isoform diversity manifests as changes in localisation. To our knowledge, this is the first time high-content imaging is used to directly examine isoform localisation at scale, and we speculate this type of technique can be used to complement the more traditional interaction-based isoform studies. Whilst relatively small, our screen identified that nearly half of tested isoform pairs differ in subcellular distribution, supporting our network-level predictions, and suggesting AS may frequently alter protein localisation. This is corroborated by a recent study by Lambourne *et al*., on protein isoforms of transcription factors that found ∼50% of isoforms show subcellular localisation differences in the form of nuclear versus cytoplasmic localisation, condensate formation, or both^36^. While Lambourne *et al.* examined complete transcript isoform changes, our analysis focused on individual splicing events and identified a similar extent of subcellular regulation. This suggests that even single exon-level variations can be sufficient to alter protein localisation, although combining multiple splicing events may produce more pronounced phenotypes. Together, this argues that considering isoform-level information is important for understanding how protein localization is specified, as well as for improving computational methods that predict protein localization from sequence^63,64^.

Our imaging data suggests localisation changes driven by AS are often graded rather than digital, fine-tuning the degree of subcellular localisation rather than completely re-localising a protein. This is illustrated in our screen by both isoforms of NUMB and ANK2, which localise to the plasma membrane but to different extents, and by the SNX2 isoforms, which localise to vesicular structures with varying enrichment. Our structural analyses indicate that such graded localisation can arise through at least two mechanisms. First, AS may insert or remove domains (or motifs)^8^. This occurs for ANK2, where alternative inclusion of a targeting module shifts the equilibrium of membrane association, yielding graded behaviour across isoforms. Second, AS event can modulate or tune existing interfaces without altering domain structure, subtly tuning molecular interactions while preserving structural integrity. This is exemplified by AS events in NUMB and SNX2, where inclusion of short exons alters existing interaction surfaces. Consistent with this, our mutagenesis of SNX2 show that differential localisation and partner specificity occur independently of the exon’s amino-acid composition, implying a structural rather than compositional mechanism. A similar mechanism operates in SRC kinase, where a microexon within the SH3 domain switches the domain’s peptide-binding preferences from a PXXP motif to a RXPXXP motif^65^, thereby fine-tuning the composition and specificity of the signalling complex. These findings support a broader model in which AS, particularly through microexons, can introduce small structural perturbations that subtly retune molecular recognition and localisation without compromising fold or protein stability.

Together, our results support a model in which alternative splicing is not a source of random variation but a conserved mechanism for precision tuning of proteome organisation by regulating protein modular architecture. By coupling interface-level adjustments to changes in localisation and interaction specificity, AS provides a flexible yet robust means of evolving new functional capacities while preserving overall cellular architecture.

## Acknowledgements

We would like to thank Benjamin J. Blencowe and Jonathan D. Ellis for sharing the protein isoform library with us, the Katarina Gaus Light Microscopy Facility at University of New South Wales for helpful advice on image acquisition and analysis, the Bioanalytical Mass Spectrometry Facility at University of New South Wales for performing mass spectrometry and Jonathan Roth for assisting with the analysis. We would also like to thank the members of the Weatheritt lab for discussions on the project and the manuscript. We further note, this research includes computations using the computational cluster Katana supported by Research Technology Services at UNSW Sydney. S.B. is the recipient of an Australian Research Council Discovery Early Career Award (DE230100271).

## Materials and methods

See tables at the end of this section for reagent (**Table 3**), plasmid (**Table 4**), and oligonucleotide details (**Table 5**). Antibody and dye dilutions are included in the text when their use is described. Figures were created using Biorender.com.

### Molecular Cloning

#### Gateway recombination

Most subcloning of protein isoform open reading frames (ORFs) was carried out using the Gateway system (Thermo-Scientific), using BP and LR clonase II enzyme mixes (donor: pDONR303, destination: mCherry-Nterminus, Cterminus-mCherry). Gateway cloning was similarly used to generate the SNX2 isoform lentiviral plasmids (Destination: pLX303^66^), and the bait and prey plasmids used for LUMIER (destinations: 3xFLAG-Nterminus or Cterminus-3xFLAG, RenillaLuciferase-Nterminus) gifted by Benjamin J. Blencowe’s group. Plasmids gifted by researchers through Addgene are listed in **Table 4**.

The Gateway BP reactions were set up according to manufacturer’s protocol. If the insert-of-interest was not in a plasmid flanked by attB1/attB2 sites, these sites were introduced by PCR with primers tailed by the attB1/attB2 recombination sites. The PCR product was then used for BP reaction. If the insert-of-interest was already in a plasmid flanked by attB1/B2 sites, the plasmid was linearised by restriction enzyme digestion prior to the BP reaction. The BP reactions were generally incubated 2-3h at +25°C before proteinase K-based inactivation, and 30% of the reaction was then used for bacterial transformation. The Gateway LR reactions were set up in a reaction volume half of that described by the manufacturer with every component halved, including DNA amounts, but where otherwise performed according to the manufacturer’s protocol. The LR reactions were generally incubated 1-2h at +25°C before proteinase K-based inactivation, and 50% of the reaction was then used for bacterial transformation. If either BP or LR reactions weren’t used immediately for bacterial transformation, the reaction was frozen at -20°C and used within a week.

#### Site-directed mutagenesis

Protein isoform ORFs not obtained from the Blencowe lab were generated by site-directed mutagenesis. Similarly, site-directed mutagenesis was used to generate the SNX2 mutants, and to introduce stop codons in a few ORFs. All site-directed mutagenesis was done using the Q5 Site-Directed mutagenesis Kit (New England Biolabs) according to the manufacturer’s protocol. All plasmids (generated via Gateway recombination or site-directed mutagenesis) were verified via Sanger sequencing at the Garvan Molecular Genetics facility.

#### Bacterial transformation

Home-made DH5α *E. coli* was used for most transformation reactions. Briefly, 50µL cells in a 2mL Eppendorf tubes were used per transformation. The cells were mixed with 1-10ng plasmid DNA, 30% of an BP reaction, 50% of an LR reaction, or 50% of a site-directed mutagenesis reaction, and then incubated on ice for 30min. The cells were then heat shocked at +42°C for 1min and then incubated on ice for 2min. 900µL room temperature LB media was then added, and the tubes were incubated in a shaking incubator (+37C, 200-250rpm) for 1h. The bacterial suspension was then plated on LB agar plates with the appropriate antibiotic for the plasmid in question. Ampicillin was used at 100µg/mL, kanamycin was used at 50µg/mL, and spectinomycin was used at 75µg/mL. Transformations of constructs that transformed less efficiently (for instance, BP reactions of large inserts), was done using commercial DH5α *E. coli* (New England Biolabs, C2987H) according to manufacturer’s protocol, using the same DNA input amounts as listed for Home-made DH5α *E. coli*. Transformation of plasmids used for lentivirus production was done using commercial OneShot Stbl3 *E. coli* (Thermo-Scientific, C737303), according to manufacturer’s protocol. The Stbl3 cells were incubated at +30°C rather than +37°C. Home-made DH5α *E. coli* was from commercial DH5α (New England Biolabs, C2987H) as previously described^67^.

#### Plasmid purification

Plasmids were purified from liquid cultures of bacterial transformants expressing the plasmid-of-interest, using various plasmid purification kits. The bacterial input amount and purification was done according to manufacturer’s protocol in all cases. Minipreps were done using Promega’s Wizard® Plus SV Miniprep kit (A1330), Midipreps were done using Macherey-Nagel’s NucleoBond Xtra Midi kit (740410), and Maxipreps were done using the Promega’s PureYield kit (A2392). Liquid bacterial cultures were grown at +37°C and 200-250rpm for non-lentiviral plasmids, and at +30°C and 200-250rpm for lentiviral plasmids.

#### Routine cell culture

HeLa cells (both wild type and stable and knockout lines) and HEK293T wild type cells were grown in non-coated TC-treated polystyrene plastic ware (Corning) in DMEM (High glucose, sodium pyruvate, Thermo-Scientific#11995065) supplemented with 10% FBS, with or without Pen/Strep (penicillin: 100 units/mL, streptomycin: 100µg/mL), at +37C, 5% CO_2_ in a humidified cell incubator. Both cell lines were passed by washing the cells 2x in heated 1xPBS (137mM NaCl, 10mM Na_2_HPO_4_, 1.8mM KH_2_PO_4_, 2.7mM KCl) then treating the cells with trypsin-EDTA 0.25% for 4min at +37°C to detach them. The cells were then resuspended in supplemented DMEM, spun at 200 x g for 4min at room temperature, and the supernatant was discarded. The cell pellet was resuspended in fresh supplemented DMEM, counted using a Countess II Automated Cell Counter, diluted further in supplemented DMEM to the desired cell density and used for downstream assays or continued culture. HeLa and HEK293T wild type cells were gifted by Ruth Lyons (Garvan Tissue Culture Facility manager).

### Cell line generation

#### CRISPR-Cas9-based gene knockout

First, we cloned gRNAs targeting SNX2, either previously described^68^ or predicted using the CRISPRscan online tool^69^, into the pSpCas9(BB)-2A-GFP (PX458)^70^ (Addgene #48138) plasmid under a U6 promoter. The gRNAs generally targeted exon junctions of exon 2 and were predicted to cause gene knockout by disrupting exon 2 usage The gRNA introduction into the plasmids was done as described by Ran *et. al.* for synthetic oligonucleotides^70^. Briefly, the gRNA oligo and the complementary sequence were ordered separately and then annealed *in vitro*. Both the gRNA and complementary strand were ordered with overhanging 5’-ends that are complementary to the BbsI-generated sticky ends in the pSpCas9(BB)-2A-GFP (PX458) plasmid. The annealed gRNA/complementary strand could therefore be inserted via into the plasmid via DNA ligation.

To generate the SNX2 KO HeLa cells, 3µg gRNA-containing pSpCas9(BB)-2A-GFP (PX458) plasmid was transiently transfected into HeLa cells in p60 dishes using ViaFect transfection reagent (4:1 reagent/DNA ratio), according to the manufacturer’s protocol. Notably, the two plasmids with different SNX2-targeting gRNAs were transfected into different p60 dishes to generate independent SNX2 KO HeLa cell lines. The cells were incubated for 48h after transfection at normal culture condition. The cells were then detached and spun as for normal cell passage, then resuspended in 1xPBS w. 2% FBS and Pen/Strep and passed through a 35µm nylon mesh to remove cell clumps. The cell suspension was then fluorescence-activated cell sorted (FACS’ed) on a BD FACS Aria III (nozzle diameter: 100µm) to isolate EGFP-positive cells positive which were sorted into 96-well plates with one cell being seeded per well. The wells contained 50µL fresh supplemented DMEM with Pen/Strep and 50µL conditioned supplemented DMEM. The single clones were then allowed to grow into individual clonal lines. Clonal lines were subsequently tested for successful gene knockout by western blotting.

Conditioned supplemented DMEM was generated by aspirating the media from HeLa cells after the cells had been incubated with the media for 24-48h. The isolated media was then spun at 3220 g x 10min at +4°C to pellet potential cell debris, and the supernatant was filtered through a 0.22µm filter. The filtered supernatant was used as conditioned supplemented DMEM.

#### Stable cell lines

We generated cell lines that stably expressed the gene-of-interest via lentiviral transduction. We generated the lentiviruses by co-transfecting 60-80% confluent HEK293T in T-25 flasks with 3 plasmids: 4µg psPAX2 (packing vector, Addgene (Plasmid #12260)), 2µg pCAG-VSVG (envelope vector, Addgene (Plasmid #35616)), and 4µg pLX303 lentiviral plasmid containing the ORF for one of the two SNX2 isoforms. The transfection was performed using Lipofectamine 3000 reagent (Thermo-Scientific) (ratios: 1.5:1 reagent/DNA, 2:1 P3000/DNA), according to manufacturer’s protocol. 24h after the transfection, the media was changed to supplemented DMEM. After another 24h, the media was isolated from the HEK293T and spun at 2500 x g for 10min at +4°C to pellet cell debris. The supernatant was then filtered through at 0.45µm filter, and this filtered supernatant was used for viral transductions (virus-containing supplemented DMEM not used immediately was frozen at - 70°C for later use).

To transduce the target cells, the virus-containing supplemented DMEM was added to the media of HeLa cells (either HeLa wild type or SNX2 knockout clones) in 6-well plates. The cells were incubated 48h with the virus and the media was then removed. The cells were washed 10x in 1xPBS, and fresh supplemented DMEM was then added, and the cells were removed from the PC2 facility. The cells were subsequently passed (after 30min of rest in a cell incubator) and seeded into new 6-well plate wells in supplemented DMEM with 2µg/mL blasticidin to select for transduced cells. The cells were treated for 4 days with blasticidin at this concentration (media changed every second day) and passed as normal when they reached confluency. After 4 days, the blasticidin concentration was increased to 10µg/mL to make sure surviving cells were transduced. This concentration was used for 6 days after which the drug was removed, and the cells were cultured as per normal to grow cells for freezing. Upon reviving frozen stocks of the stable cells, they were cultured at 3µg/mL blasticidin when not used for experiments to keep selecting for cells expressing the transgene. The drug was removed when cells were used for AP-MS or immunofluorescence. Expression of the transgenic SNX2 (either +EXON or –EXON isoforms) was verified by western blotting and end-point RT-PCR.

Generation of stable cells by viral transduction was done in a Garvan PC2 laboratory designated for viral work (the point at which the cells were removed from the facility is specified). This was done in accordance with Garvan’s PC2 guidelines.

### Cell preparation for imaging

#### Protein isoform library

HeLa wild type cells were detached, counted, and diluted in supplemented DMEM (as above) to make a cell suspension with a density of 90k/mL. 500µL of the cell suspension was then added into each well in a glass-bottom 24-well plate (Cellvis) (45k cells/well). Immediately after adding the cell suspension to the wells, the cells were reverse transfected by dripping in 50µL transfection mix consisting of 400ng plasmid encoding mCherry-tagged construct mixed with ViaFect (4:1 reagent/DNA ratio) in OPTI-MEM as per manufacturer’s protocol. Each well was transfected with a different construct (except the negative controls). One well was left untransfected, and instead 50µL OPTI-MEM was added. Three experimental replicates for a given protein isoform batch were set up sequentially on the same day with cell suspensions and transfection mixes being made separately for each replicate.

After seeding and reverse transfection, the cells were incubated for 24h in the cell incubator. After the incubation, all wells were washed 2x in 500µL heated 1xPBS and then fixed with 200µL 4% paraformaldehyde for 10min at room temperature. The wells were subsequently washed 3x in 500µL 1xPBS, and 1mL 1xPBS was then added to each well. The well plate was then wrapped in parafilm and kept at +4°C until used for immunofluorescence staining (within 7 days of fixation).

#### SNX2 mutants

HeLa wild type cells were detached, counted, and diluted in supplemented DMEM (as above) to make a cell suspension with a density of 100k/mL. 450µL of the cell suspension was then added into each well in a poly-D-lysine coated glass-bottom 24-well plate (Cellvis) (45k cells/well). Cells were incubated for 16h in a cell incubator and were then forward transfected with plasmids encoding mCherry-SNX2 constructs (both endogenous isoforms and mutants) or mCherry itself by dripping in 50µL transfection mix consisting of 375ng plasmid DNA mixed with jetPRIME (2:1 reagent/DNA ratio) in jetPRIME buffer as per manufacturer’s protocol. Each well was transfected with a different construct (except 3 wells that were all transfected with mCherry-SNX2 -EX). One well was left untransfected, and instead 50µL jetPRIME buffer was added. Three experimental replicates were set up sequentially on the same day with cell suspensions and transfection mixes being made separately for each replicate.

The cells were fixed 24h after transfection by removing the media and adding 200µL 4% paraformaldehyde for 10min at room temperature. The fixed sample was washed and stored as for the protein isoform library samples.

#### Untagged SNX2 in HeLa SNX2 cell lines

HeLa wild type and SNX2 lines (knockout and rescue cells) were handled as described for ‘SNX2 mutants’ to generate samples for imaging of either untagged, stably expressed SNX2 isoforms (non-transfected wells) or mCherry-SNX2 isoforms in the knockout background.

#### Immunofluorescence staining of protein isoform pairs and SNX2 mutants

The well plates with fixed, transfected cells were stained for imaging within 7 days of fixation. All replicates for a given protein isoform batch or SNX2 mutants were stained at the same time. The staining was done as follows: (note, between each step, the wells were washed 2-4x with 500µL 1xPBS per wash. During incubations, the well plates were covered with alu foil). 1) Permeabilisation. Cells were permeabilised in 200µL 1xPBS with 0.2% Triton-X100 for 12min at room temperature. 2) Blocking. The cells were then blocked in 200µL 1xPBS with 3% BSA for 60min at room temperature. 3) Primary antibody staining. The cells were then stained in 200µL 1xPBS with 3% BSA and mouse anti-Golgin97 (0.4µg/mL) for 120min at +37°C. 4) Secondary antibody staining. The cells were then stained in 200µL 1x PBS with 3% BSA and goat anti-mouse-Alexa488 (3µg/mL) for 60min at room temperature. 5) DAPI staining. The cells were then stained in 200µL 1xPBS with DAPI (1µg/mL) for 15min at room temperature. 6) NHS-ester staining. The cells were then stained in 200µL 1xPBS with NHS ester-Alexa647 (0.125µg/mL) for 30min at room temperature (note, the NHS ester was made up just before being added to the cells). 7) Final wash. The cells were washed 4x in 500µL 1x PBS and a final 1mL of 1xPBS was added to each well to prevent the samples from drying out. The stained well plates were wrapped in parafilm and kept at +4°C until used for imaging (typically less than a week).

#### Immunofluorescence staining of SNX2 in HeLa SNX2 lines

The samples were stained as described in ‘Immunofluorescence staining of protein isoform pairs and SNX2 mutants’, except the primary antibody was a mouse anti-SNX2 antibody (1.25µg/mL).

### Imaging

#### Imaging of protein isoform library

Fixed and immunofluorescence-stained cells were imaged using a PerkinElmer Operetta CLS system using its spinning disk confocal setting. The cells were imaged with a 40x water immersion 1.1NA objective, resulting in fields-of-view of 0.32mm x 0.32mm (2160 x 2160 pixels), with a pixel size of 0.148µm x 0.148µm. We acquired ∼50 fields-of-view (FOV) per sample per experimental repeat with 14 slices being acquired per FOV with 0.5µm between each slice (z-dimension). Three biological repeats were generated for each isoform batch. Four channels (referred to as DAPI, Golgin97-Alexa488, mCherry, and NHS Ester-Alexa647) were acquired for each slice using the following excitation, emission, and exposure settings (**Table 1**). Note, the microscope uses LEDs with bandpass filters (BPF) rather than lasers for excitation.

**Table 1.**
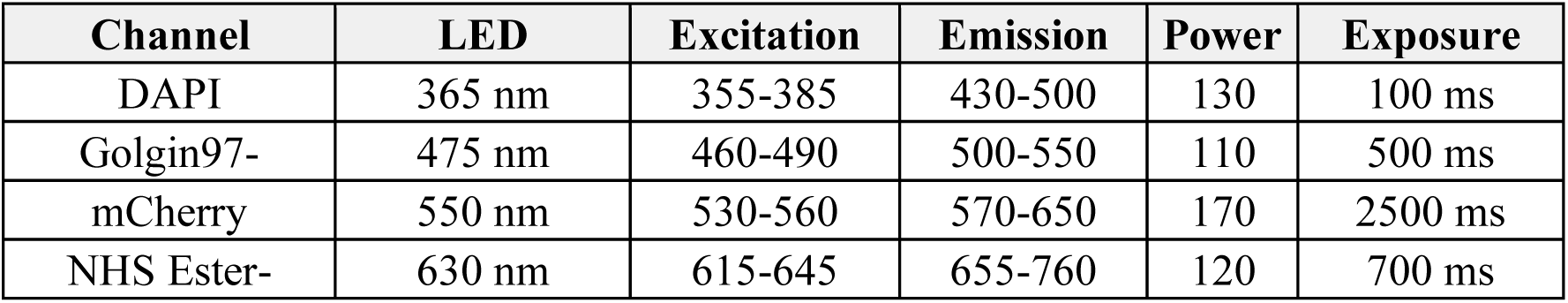
Excitation, emission, and exposure for the four acquired channels.

| Channel | LED | Excitation | Emission | Power | Exposure |
| --- | --- | --- | --- | --- | --- |
| DAPI | 365 nm | 355-385 | 430-500 | 130 | 100 ms |
| Golgin97- | 475 nm | 460-490 | 500-550 | 110 | 500 ms |
| mCherry | 550 nm | 530-560 | 570-650 | 170 | 2500 ms |
| NHS Ester- | 630 nm | 615-645 | 655-760 | 120 | 700 ms |

#### Imaging SNX2 mutants

The SNX2 mutant samples were imaged similarly to the protein isoform library except only 8 slices were acquired per FOV with a step size of 1.0µm. 41 FOVs were acquired per sample per repeat. Three biological repeats were generated.

#### Imaging of SNX2 in HeLa SNX2 lines

The HeLa SNX2 cell line samples were imaged similarly to the protein isoform library except only 8 slices were acquired per FOV with a step size of 1.0µm. 46 FOVs were acquired per sample per repeat. Here, the green channel corresponded to SNX2. Three biological repeats were generated.

#### Image processing, cell measurements, cell filtering, and transfection status Protein isoform screen and SNX2 mutants

We performed analysis of single slices from the acquired confocal stacks, using the slice corresponding to where the cell adheres to the glass surface.

The acquired confocal stacks were processed as outlined in **Figure S2**. This was done by: **1)** identifying the focus slice in each confocal stack using ImageJ v1.54f^71^ and the plugin Select focus slice v1.0 (unpublished, from: https://sites.google.com/site/qingzongtseng/find-focus, downloaded June 2022) with a user-defined variance threshold on the NHS-Alexa647 channel. The same threshold was applied to all images for a given repeat. The selected focus slice was then used for all channels for the given FOV. **2)** We illumination corrected all channels in ImageJ using the BaSiC v1.0 plugin^72^ where all focus images from the experimental repeat for the given channel were used to calculate the correction profiles (both flat and dark field) using default settings. The correction profiles were then used to illumination correct the images. **3)** We segmented individual cells using CellPose 2.0^73^, using the illumination corrected NHS-Alexa647 and DAPI as input to define the cell body, and nucleus, respectively. We used a custom segmentation model for segmentation generated via transfer learning using the default ‘cyto’ model with default settings as the starting model. Cell diameter was set to 150 pixels. **4)** We then trained a custom classifier to label cells as either interphase or mitotic using Ilastik v1.4^74^, based on cell morphology and NHS staining, to focus analysis on interphase cells. **5)** We then loaded the illumination corrected channel images and cell segmentation maps into CellProfiler v4.1^75^ and first subtracted the median background (defined as the part of the image not covered by cells) from all channel images. **6)** We then measured features describing cell morphology, the mCherry channel, and the Golgin97 channel, as well as correlation between Golgin97 and mCherry. **7)** The dataset was then filtered to remove mitotic cells, cells on image borders, non-mononucleated cells, cells with touching nuclei (typically a result of mis-segmentation of multinucleated cell into two cells), and cells where >85% of their cell border contacts other cells as this typically indicated tight packing of cells, which impaired segmentation accuracy. **8)** We then assigned transfection status to the cells based on their cell body mean mCherry intensity and 75% quantile mCherry intensity, where both values had to surpass user-defined thresholds based on the non-transfected cell sample. **9)** For transfected cells, we removed the brightest cells from the transfected populations on a pairwise-basis that reflected which samples would be compared. We defined an outlier threshold for each sample as: 𝑂𝑢𝑡𝑙𝑖𝑒𝑟 = ^𝑚𝐶ℎ𝑒𝑟𝑟𝑦^𝑀𝑒𝑎𝑛𝐼𝑛𝑡𝑒𝑛𝑠𝑖𝑡𝑦 ^> 𝑃𝑜𝑝𝑢𝑙𝑎𝑡𝑖𝑜𝑛 𝑚𝐶ℎ𝑒𝑟𝑟𝑦^75%𝑄𝑢𝑎𝑛𝑡𝑖𝑙𝑒𝐼𝑛𝑡𝑒𝑛𝑠𝑖𝑡𝑦 ^+ 1.5 ⋅𝑃𝑜𝑝𝑢𝑙𝑎𝑡𝑖𝑜𝑛 𝑚𝐶ℎ𝑒𝑟𝑟𝑦^𝐼𝑛𝑡𝑒𝑟𝑄𝑢𝑎𝑛𝑡𝑖𝑙𝑒𝑅𝑎𝑛𝑔𝑒

We then applied the lower threshold for a given isoform to both samples in that pair. Notably, this filtering step generally removed very few cells and so only removed the brightest cells. Step 9) was not performed for the SNX2 mutant experiment as we didn’t perform pairwise comparisons. Instead, we removed the outliers for each population using the outlier threshold calculated (as above) for the same population.

The non-transfected and transfected cells made up the cells used for downstream analysis.

#### SNX2 cell lines

We analysed the images as described in ‘Protein isoform screen and SNX2 mutants‘, but also identified whether the cells expressed SNX2 or not (similar to step 8 used to determine mCherry transfection status). Here, the wells stained without a primary antibody was used to define the intensity thresholds (mean SNX2-Alexa488 intensity and 75% quantile SNX2-Alexa488 intensity) to determine cells expressing and not expressing SNX2. Notably, this was necessary, as not all cells in the Rescue lines were expressing SNX2.

#### Representative images

Representative images were selected based on our analyses of the overall population behaviour of the given mCherry constructs (for isoform imaging and SNX2 mutants) or SNX2 isoforms (for SNX2 cell lines). These images were prepared in ImageJ v1.54f using the illumination corrected channel images. All images that are directly compared have the same contrast settings. Mean mCherry intensities for representative cells were determined by manually segmenting the cells and measuring their mean mCherry intensity, then subtracting the mean background mCherry intensity measured adjacent to each representative cell. The values were then normalised to the transfection threshold, giving the mCherry construct amount relative to the effective detection limit. These values directly correspond to the plotted intensity values in the population XY plots. The same approach was used to determine SNX2 expression levels in the SNX2 cell lines, although the expression value was normalised to the median intensity observed in the HeLa wild type population rather than the transfection threshold, which gave us the SNX2 abundance relative to the endogenous level.

### Analysis of cell measurements

#### Classifier-based analysis of protein isoform pairs

We used the single cell feature measurements to compare localisation of mCherry constructs (such as two isoforms from the same gene) in a pairwise-manner using a classifier-based approach as outlined in **Figure S3**. We performed separate analyses for transfected and non-transfected cells from the same wells. These made up the ‘isoform’ and ‘internal control’ groups, respectively.

The analysis consisted of 1) generating an appropriate dataset through sampling, and 2) analysing the dataset through a classifier-based approach. For 1), we pooled transfected cells expressing the two isoforms to be compared across replicates, then down-sampled the cells in the more abundant isoform group to match the number of cells of the less abundant isoform in a replicate-aware manner. Accordingly, replicates could contribute different numbers of cells compared to each other, but the cells contributed by a replicate was always 50% isoform A and 50% isoform B. Cell numbers used for classifier assessment after down-sampling and pooling was ≥170, although most pairs had substantially more cells (median cell number = 967). Cell numbers for each comparison are listed in **Table S3** The dataset from 1) was then used for classifier-based analysis 2), where the classifier was trained to assign the correct isoform identity (corresponding to well identity) based on the measured features.

The classifier-based analysis was performed in R (4.3.3) using Caret v6.0-94^76^ to control classifier training, testing, and feature assessment, with the classifier itself being a feedforward neural network, from nnet^77^ v7.3-19. The initial input features for the classifier were all features describing the mCherry signal (although intensity features were normalised to mean mCherry intensity) and cell morphology. Zero variance features were then removed, and the remaining features (around 200 for all classifiers) were centred, then scaled, before being used for classifier assessment. For the assessment, the classifier was trained with 10-fold cross-validation, and we used the average ROC-AUC (Receiver Operating Characteristic (ROC) Area Under the Curve (AUC)) at the parameters resulting in the best performance, across the test folds as the metric describing the degree of isoform differences. The 1) sampling and 2) classifier assessment was repeated 50 times.

To normalise the ROC-AUC metric for potential well-specific effects driving classification, we performed the same analysis on non-transfected cells from the same wells as the two mCherry constructs being compared and used these as internal controls. Notably, the internal controls were sampled to best mirror the isoforms in question, and the non-transfected cells were therefore sampled to match the proportion of cells contributed by each replicate for the transfected cells, while also contributing the same number of cells from the two wells.

Normalised ROC-AUCs were calculated using the ROC-AUC from the ‘Isoform’ and ‘Internal control’ classifiers as outlined in **Table 2** (same as **Figure S4C**):

**Table 2.**
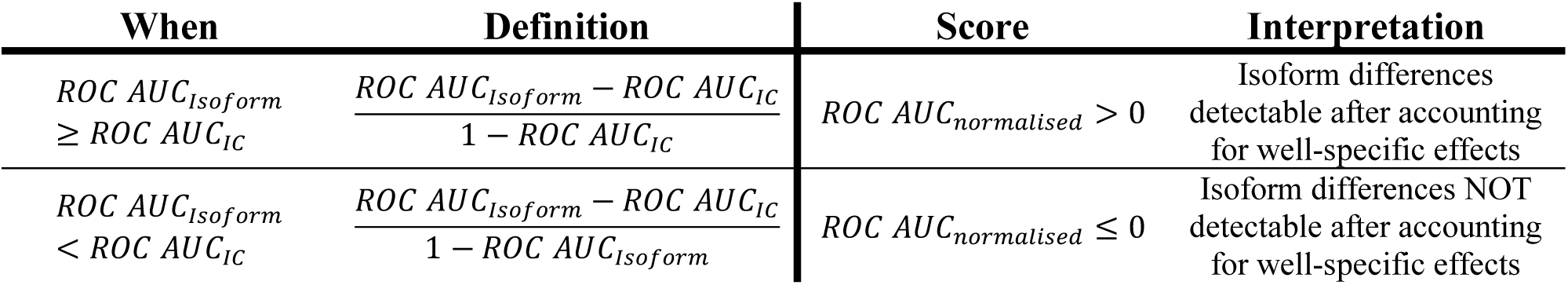
Definition and interpretation of normalised ROC-AUC performance metric.

The normalised ROC-AUC ranges from -1 and 1 and is symmetric around 0. Scores above 0 indicates mCherry construct differences can be detected after accounting for well-specific effects.

We statistically tested whether the normalised ROC-AUC for a given isoform comparison was significantly greater than zero using a one-sided bootstrap approach in R using boot v1.3-31^78^. For each pairwise comparison, we performed 2000 bootstrap iterations by resampling with replacement from the normalised ROC-AUC values, calculating bootstrap confidence intervals for the mean performance. If zero was excluded from the lower bound of the 99% one-sided confidence interval, the isoform classifier was determined to significantly outperform the internal control classifier, indicating isoform differences. We corrected for multiple testing using the Benjamini-Hochberg method to control the false discovery rate.

#### Classifier-based analysis after intensity binning

We examined whether our classifier-based analysis yielded consistent results across the mCherry expression range for isoform pairs by separating transfected cells for a given isoform pair into lower and higher intensity groups and then perform the classifier-based analysis as described in ‘Classifier-based analysis of protein isoform pairs’ separately on the groups. Cells were grouped using two separate approaches (App. 1 or App. 2) as outlined in **Figure S9A**. In approach 1, we identified the max single cell mean mCherry intensity for both isoforms in a pair, selected the lower of the two, and assigned cells expressing either isoform into bins as: lower bin = mean mCherry intensity < 20% of max. Higher bin = mean mCherry intensity > 20% of max. In approach 2, we identified the median single cell mean mCherry intensity for both isoforms in a pair, selected the lower of the two, and assigned cells expressing either isoform into to bins as: lower bin = mean mCherry intensity < median. Higher bin = mean mCherry intensity > median.

We compared the classifier results for all isoform pairs with and without binning by plotting the normalised ROC-AUC in R (4.3.3) using pheatmap v1.0.12^79^. We compared the population mCherry level for all isoform pairs with and without binning by plotting the population median of mean mCherry intensities, relative to transfection threshold.

#### Out-of-sample testing

We assessed out-of-sample performance of classifiers trained on controls and select isoform pairs by using cells from one batch (batch A or B) for the training set, and cells from the other batch for testing. For the training set, we sampled and trained the classifier as described in ‘Classifier-based analysis of protein isoform pairs’. For the test set, we sampled the cells as described in ‘Classifier-based analysis of protein isoform pairs’, to have the same number of cells of the two groups and then predicted isoform identity for the cells using Caret^76^ predict(). The performance was assessed as accuracy. Notably, we sampled the test set to have equal classes to both mirror the training set and to make the accuracy comparable between conditions (as accuracy is sensitive to class imbalance). The accuracy can therefore range from 0.5 to 1.0. 20 iterations of classifier training and testing was done, and the performance values were the mean accuracy from each iteration. For each iteration, a classifier was trained on the training batch and was then used to predict isoform identities for the down-sampled test batch, where the test batch down-sampling and prediction was performed 50 times. The mean accuracy across the 50 times was used as the performance.

To examine the association between the accuracy of the out-of-sample tests and the normalised ROC-AUC approach, we performed linear regression, Accuracy ∼ normalised ROC-AUC. We statistically tested if the accuracy and normalised ROC-AUC were correlated in R using a correlation test, cor(method = ‘kendall’).

#### Replicate sampling imbalance

We quantified replicate sampling imbalance as the maximum fraction of cells used in given isoform pair comparison that came from a single experimental repeat over the total number of cells used for the comparison (cells pooled across all repeats).

#### mCherry intensity imbalance

We quantified mCherry intensity (expression) imbalance between isoforms in a pair by first calculating the population median of single cell mean mCherry intensities for each isoform of the cells used for classifier analysis. We then determined the population difference as: 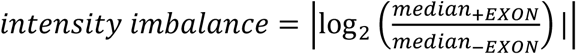. This was calculated for each classifier iteration as cell sampling was part of each classifier iteration, and the final isoform imbalance score was defined as the median of the intensity imbalance scores for the given isoform pair.

#### Association between imbalance and performance

The correlation between normalised ROC-AUC and imbalance metrics (replicate sampling imbalance, mCherry intensity imbalance) was statistically examined in R using a correlation test, cor(method = ‘kendall’).

#### Identifying Isoform distinguishing features

Using the default classifier importance scores generated by Caret for the nnet classifiers, we first calculated the Feature contribution percentage (%), that specified how large a percentage of the sum of feature importance scores were contributed by a given feature:

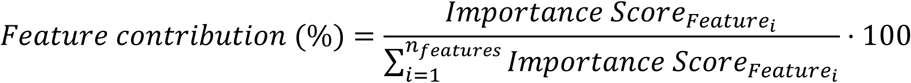

(n features: number of features used for the given classifier, ∼200 for all classifiers).

We did this for both the isoform classifier and associated internal control, which allowed us to calculate contribution ratios, specifying the relative contribution of a feature to the isoform classifier compared to the internal control:

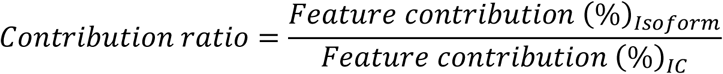

(Isoform: isoform, IC: internal control).

We then defined isoform distinguishing features (features that were notably more important for the isoform classifier compared to the associated internal control classifier), as features with a contribution ratio and a contribution ratio multiplied with the feature contribution percentage for the isoform classifier above two user-defined threshold values.

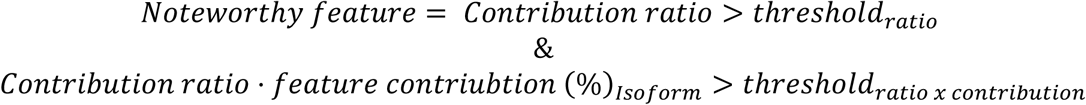

The user-defined threshold values were defined so the negative control had no distinguishing features. The contribution ratio threshold was 2.6, and the ratio x contribution threshold was 1.5.

#### Principal component analysis of SNX2 mutants

We analysed the SNX2 mutant constructs by principal component analysis (PCA) in R. As input, we used the mean values of the distinguishing features identified for the two SNX2 isoforms (about 20), after highly correlated features had been removed using findCorrelation(cutoff=0.8, exact=TRUE) from Caret v6.0-94^76^. The mean values of the distinguishing features were calculated across the entire cell population (pooled across the three repeats) for the given construct, with the lowest number of cells for a construct being n = 185, while most constructs had n > 300 (cell numbers for each construct listed in **Figure S13B**). The feature means were centred and scaled before performing PCA using prcomp(). The plots of PCA scores, biplots, and the scree plots were created using the factoextra^80^ and tidyverse^81^ packages. We performed this analysis separately for transfected and non-transfected cells.

#### Affinity purification mass spectrometry

We examined the protein interactomes of the SNX2 isoforms using affinity purification mass spectrometry (AP-MS) on HeLa wild type and HeLa SNX2 cell lines. The immunoprecipitation protocol was based on^82^ with minor modifications. To isolate SNX2 and its interactors, we grew HeLa wild type and SNX2 rescue cells lines to 80-90% confluency in P150 dishes in supplemented DMEM with Pen/Strep. The cells were then washed 2x in ice cold 1xPBS, after which 1mL 1xPBS was added to each dish and the cells were detached by cell scraping. The detached cells were transferred to Eppendorf tubes and spun at 800 x g for 5min at +4°C to pellet the cells. The supernatant was then removed, and the cell mass was weighed and then snap frozen in liquid nitrogen. The cell pellets were stored at -70°C until the immunoprecipitation was continued (within a week of harvest). To continue the immunoprecipitation, the cell pellets were thawed on ice and then resuspended in immunoprecipitation buffer (20 mM HEPES pH 7.2, 100 mM K-acetate, 2.5 mM Mg-acetate, 0.1% Nonidet-P40 (NP40) (v/v), 1mM DTT, 10% glycerol (v/v)) at a ratio of 4µL/mg pellet by gentle pipetting. The cell suspensions were then frozen on dry ice and thawed by holding the tubes to promote cell lysis (freeze-thaw). The lysates were then transferred to a tube spinner and spun at 12rpm for 30min at +4°C. The cell lysates were then transferred to a 2mL Dounce homogenizer for mechanical shearing (25 strokes). Homogenised lysates were then transferred to Eppendorf tubes and spun at 17000 x g for 20min at +4°C to pellet nuclei. The supernatant was recovered and transferred to a new Eppendorf tube. 15µL of the isolated supernatant was transferred to a separate Eppendorf tube and served as a measure of the input fraction for western blotting (kept at -20°C until use). The remaining supernatant was mixed with 5µL anti-SNX2 antibody (corresponding to 1.25µg antibody) and incubated on a tube spinner at 7rpm overnight at +4°C. The next day, the antibody-lysate mix was mixed with 50µL immunoprecipitation buffer-equilibrated Protein G Dynabeads or Protein A Dynabeads (Thermo-Scientific), and the bead lysate suspension was then incubated on a tube spinner at 7rpm for 2h at +4°C. After this incubation, the suspension was put on a magnetic rack to pull down the beads and the antigen-antibody complexes. The supernatant was removed and transferred to a separate tube that was frozen at -20°C (it served as the unbound fraction for western blotting). The beads were then resuspended in 1mL immunoprecipitation buffer and transferred to a new tube. The tubes were then placed on a magnet again and the supernatant was removed. The beads where then resuspended in 1mL wash buffer (20 mM HEPES pH 7.2, 10mM MgCl_2_) by gentle pipetting and placed on the magnet a final time and the supernatant was removed. The sample was then eluted from the beads by adding 30µL 4xLDS sample buffer diluted to 1x in 1xPBS with 20mM DTT. The sample was then incubated at +95°C for 5min, after which it was put back on magnet so the eluate could be isolated from the beads. 5µL of the eluate was kept as the immunoprecipitated fraction for western blotting and the remaining 25µL was used for mass spectrometry analysis. Eluted samples were kept at -20°C until use (within a week). Samples were subjected to tryptic digest (bottom-up) prior to mass spectrometry analysis on a Thermo-Scientific LTQ Orbitrap XL instrument. Downstream sample processing, including tryptic digest, and label-free quantification mass spectrometry analysis was performed by the Bioanalytical Mass Spectrometry Facility at University of New South Wales. The mass spectrometry data was analysed using SAINT^83^ using default settings. The experiment was repeated three times.

#### LUMIER

Luminescence-based mammalian interactome mapping (LUMIER)^84^ was performed to examine binary protein-protein interactions as described^46^, with slight modifications. Briefly, HEK293T wild type cells were passaged (see ‘Routine cell culture’) and seeded into wells in a 48-well plate (TC-treated, non-coated polystyrene) at 40k cells/well in 200µL supplemented DMEM. The cells were incubated for 24h and then co-transfected with 200ng total plasmid DNA (100ng prey plasmid, and 100ng bait plasmid) for the various interactions tested. The transfection was done using Lipofectamine 3000 (ratios: 1.5:1 reagent/DNA, 2:1 P3000/DNA) with 20µL transfection mixture made up in OPTI-MEM being dripped into each well and mixed with the media by gentle pipetting. Every transfection combination was done in triplicates in each experiment, and these wells were subsequently handled in parallel and were used as technical replicates (observations from same experiment repeat). The cells were incubated for 48h with the transfection media, and each well was then washed in 1x in 1xPBS and lysed in 150µL LUMIER lysis buffer (50 mM Tris–HCl pH 7.4, 150 mM NaCl, 1 mM tetrasodium EDTA, 0.5 % Triton-X100 (v/v), and Halt protease/phosphatase inhibitor. Halt added right before use). The wells were incubated with the lysis buffer for 40min at +4°C under constant, gentle agitation on an orbital shaker. 90µL of each lysate was then transferred to a Lumitrac 96-well plate precoated with anti-FLAG antibody (described below). This served as the IP plate. Another 20µL of each lysate was transferred to an uncoated Lumitrac 96-well plate. This served as the TOTALs plate. Both plates were sealed with sealing tape and incubated for 60min at +4°C without agitation. The lysate was then removed from the IP plates by aspiration and the IP plate was washed 6x in 250µL LUMIER wash buffer (50 mM Tris–HCl pH 7.4, 150 mM NaCl, 1 mM tetrasodium EDTA, and 0.1 % Triton-X100 (v/v)). The wash buffer was then removed and 100µL Renilla-GLO reagent (Promega) (made fresh according to manufacturer’s instructions) was added to each well. Right after, 20µL Renilla-GLO reagent was added to each well in the TOTALs plate. Both plates were incubated for 10min at RT with the reagent, and the luminescence was then read on a CLARIOstar plate reader (BMG). The IP plate was then used to measure the amount of prey protein by ELISA as follows: Each well was washed 8x in 250µL PBS-T (1x PBS, 0.05% Tween 20 (v/v)). The PBS-T was then removed and 100µL LUMIER anti-FLAG-HRP antibody solution (1x PBS, 1% FBS (v/v), 5% Tween 20 (v/v) with anti-FLAG-HRP (0.05µg/mL). Antibody added fresh) was added to each well. The plate was then sealed and incubated for 30min at +4°C without agitation. The wells were then washed another 8x in 250µL PBS-T. The PBS-T was removed and 100µL ELISA Pico substrate (made fresh) was added to each well. The plate was incubated for 7min at RT, and the absorbance was then read on a CLARIOstar plate reader (BMG). The experiment was repeated 3-4 times for all examined interactions.

#### LUMIER data analysis

The LUMIER interaction scores, NLIR and NtFLIR, were calculated using luminescence measurements of the TOTALs and IP plate and IP ELISA as described^46^. NLIR describes luminescence signal in the IP plate divided by luminescence in the TOTAL plate. NLIR > 3 were considered positive interactions^46^. NtFLIR was calculated as the NLIR score divided by the absorbance measured in the ELISA against FLAG (which quantifies Prey amount) and served as an interaction score (NLIR) normalised to the amount of prey protein present.

For each experimental repeat, we used the mean NLIR and NtFLIR across the technical replicates. We calculated the interaction differences between SNX2 isoforms with a given prey protein as: 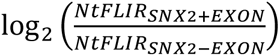, which we calculated for each experimental repeat. The SNX2 -EX isoform was used as the reference for the SNX2 +EX isoform as well as for all SNX2 mutants, where interaction differences were calculated in the same way. We statistically tested if the SNX2 +EX isoform or SNX2 mutants interacted differently with prey proteins compared to SNX2 -EX using two-sided one-sample t-tests on the interaction difference score, where the interaction differences for the different SNX2 constructs were compared to zero. We corrected for multiple testing using the Benjamini-Hochberg method to control the false discovery rate.

#### Anti-FLAG plate coating

Lumitrac 96-well plates were coated with anti-FLAG antibody by incubating each well with 100µL 1x PBS with anti-FLAG antibody (10µg/mL) overnight at +4°C. The next day, the antibody solution was removed and 250µL LUMIER blocking buffer (1xPBS, 3% BSA (w/v), 5% sucrose (w/v), and 0.5% Tween 20 (v/v)) was added to each well and incubated at 60min at RT at gentle agitation on orbital shaker. The blocking buffer was then removed, and the plate was sealed and kept at +4°C until use.

#### Western blots

Western blots were performed using standard protocols. Briefly, all protein samples were run on Bolt Bis-Tris 8% gels (Invitrogen) using 10-30µg input protein per sample (determined through a Bradford assay according to manufacturer’s protocol) or, for immunoprecipitated samples, a specific percentage of the isolated fractions for (1/150 input/unbound and 1/3 immunoprecipitated sample). After gel electrophoresis, the proteins were transferred to methanol-activated PVDF membranes (Immobilon). The membranes were then blocked with TBS-T (1x TBS, 0.1% Tween 20 (v/v)) and 6% skim milk powder (w/v) for 1h under constant, gentle agitation on orbital shaker. The membranes were then briefly washed in TBS-T, and the membranes were then transferred to small plastic bags and covered with primary antibody solution, consisting of TBS-T, 1% BSA (w/v), and the primary antibody (see below for concentrations). Note, blots were typically cut into pieces to probe different parts of the membrane for specific proteins (based on size) after blocking but prior to primary antibody incubation. The membranes were incubated with the primary antibody solution at +4°C overnight under constant agitation. The next day, the membranes were washed 3x for 10min at room temperature in TBS-T under constant gentle agitation. Then the membranes were incubated with HRP-conjugated secondary antibody solution, consisting of TBS-T, 1% skim milk powder, and secondary antibody-HRP raised against the host species of the primary antibody (see specifics below), for 1h at room temperature under constant agitation. This was also done in plastic bags. The membranes were then washed again for 3x10min at room temperature with TBS-T under constant gentle agitation. The membranes were then developed with Clarity Western ECL Substrate (BioRad) according to the manufacturer’s instructions and imaged on a ChemiDoc.

To quantify the amount of SNX2 protein, the intensity of all bands-of-interest were measured in ImageJ v1.54, after which the background was subtracted. The amount was then normalised to the amount of GAPDH (reference gene) from the same sample, measured in the same way SNX2.

#### SNX2 cell line verification blots

To verify knockout or transgene expression, cells from HeLa wild type, SNX2 KO, and SNX2 rescue cell lines were washed in 1xPBS then lysed in RIPA buffer (150mM NaCl, 1% Nonylphenol ethoxylated (v/v), 0.5% Sodium deoxycholate (w/v), 0.1% SDS (w/v), 17.1mM EDTA, 50mM Tris(HCl), pH 8) on ice for 5min. The lysates where then transferred to Eppendorf tubes, spun at 17000 x g for 10min at +4C, and the supernatant was isolated and used for western blots. Cell lysates were mixed with LDS sample buffer (Thermo-Scientific) and DTT for a final concentration of 1x, and 20mM, respectively, and examined via western blotting as described above. The primary antibodies were mouse anti-SNX2 used at 0.25µg/mL (1/1000), and mouse anti-GAPDH used at 0.04µg/mL (1/5000). The secondary antibody was goat anti-mouse-HRP, used at 0.16µg/mL (1/5000). The experiment was performed once.

#### SNX2 immunoprecipitation verification blots

The three isolated fractions (input, unbound, and immunoprecipitated) were diluted in 1x PBS; for the input fraction, and the unbound fraction this was 1/150, and for the immunoprecipitated fraction, this was 1/3. The diluted samples were then prepared in 4x LDS sample buffer and 20mM DTT and examined via western blotting as described above. The primary antibodies were mouse anti-SNX2 used at 0.25µg/mL (1/1000), and mouse anti-GAPDH used at 0.04µg/mL (1/5000). The secondary antibody was goat anti-mouse-HRP, used at 0.16µg/mL (1/5000). The experiment was performed once.

#### End-point RT-PCR

End-point reverse transcription polymerase chain reaction (RT-PCR) was carried out to examine inclusion of SNX2’s microexon to verify correct isoform expression in HeLa SNX2 stable cell lines. We isolated RNA for the analysis from the HeLa SNX2 cell lines by washing adherent cells 2x in 1xPBS, then lysing them in TRIzol reagent (Thermo-Scientific). Total RNA was subsequently purified according to the manufacturer’s protocol for TRIzol-based RNA extraction. The precipitated RNA was redissolved in RNase-free H_2_O. The reverse transcription and subsequent PCR amplification was done as a single step using QIAGEN OneStep RT-PCR kit. The reactions were set up in 10µL total volume, as described^85^. Briefly, each 10µL reaction consisted of: 3.7µL RNase-free H_2_O, 2mL 5x QIAGEN OneStep RT-PCR Buffer, 1.2µL 5µM FORWARD PRIMER, 1.2µL 5µm REVERSE PRIMER, 0.4µL 10mM dNTP Mix, 0.5µL enzyme mix, 1µL RNA template (10-30ng/µL) in RNase-free H_2_O. The reactions were run in a Thermocycler with the following steps. 1) Reverse transcription: 30min at +50C, 2) Initial PCR activation: 15min at +95C, 3) 3-step cycling (40 cycles): Denaturation, 1min at +94°C. Annealing, 1min at +54°C. Extension, 1min at +72°C. 4) Final extension: 10min at +72°C. The reactions were loaded on 2% agarose gels containing GelRed (Biotium), separated via gel electrophoresis, and imaged on a ChemiDoc to examine RT-PCR product sizes. Size differences corresponding to the size of the microexon indicated which isoform was expressed. The primer sequences for the primers used to examine SNX2’s microexon are listed in **Table 5**. The experiment was performed once.

#### Correlation between normalised ROC-AUC and LUMIER interactions

We examined the correlation between normalised ROC-AUC (localisation readout) and the overall PPI differences as quantified by LUMIER for isoform pairs where ≤3 interactions had been tested across ≤2 experimental repeats and resulted in positive interactions with at least one isoform of the pair. The LUMIER data originated from several publications by the same authors^13,23,35^ and can be found in **Table S6**.

We calculated overall PPI differences between two isoforms from the LUMIER data by calculating the fold-change in NLIR (NLIR_FC_) between isoforms for a given interactor for each repeat:

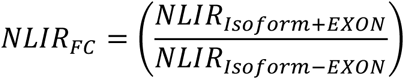

Then averaging the NLIR_FC_ for all experimental repeats and calculating the absolute log_2_ fold-change for the averaged value:

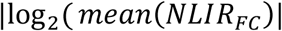

This score was calculated for each interactor that had been examined for an isoform pair, and we used the mean of these interaction difference scores as a measure of the overall PPI difference for an isoform pair. For the localisation readout, we used the median normalised ROC-AUC. We statistically tested the correlation between the median normalised ROC-AUC and collective LUMIER interaction score in R using a correlation test, cor(method = ‘kendall’). A p-value ≤ 0.05 was considered significant.

#### Analysis of microexons in α-helices

We extracted microexons (nucleotide (nt) length <30 nt) from vastDB^24^ v11 and filtered them to only include alternatively spliced microexons showing tissue-specific regulation. We specifically used microexons that were differentially included in neural tissues compared to non-neural tissues (using vast-tools compare), and with a percent spliced in (PSI) greater >15 in neurally annotated adult tissues in vastDB resource. This captures most tissue-regulated microexons^23^.

The localization of microexons within secondary protein structures was assessed using DSSP (database of secondary structure) assignments based on protein entities in the Protein Data Bank (PDB)^86^. Here, we defined α-helices as annotated by DSSP as H (α-helix) or G (3_10_-helix), β-strands as B (isolated β-bridge) or E (extended β-strand), loops as S (bend) or T (hydrogen-bonded turn). Microexons were considered within a secondary structure element if any of its residues (or the immediately preceding or proceeding residue) was annotated as a secondary structure element by DSSP. The relative position of microexon within α-helices was defined based on the minimum distance from microexon to the beginning or end of the helix as defined by DSSP.

We analysed the residue composition of microexons, and flanking residues compared to those of larger cassette exons (≥30nt), that followed a similar inclusion pattern to that of the microexons (see above). Only exon lengths divisible by 3 were included. Residues were grouped as follows: Aliphatic residues: Ile (I), Leu (L) or Val (V). Charged residues: Glu (E), Lys (K), Asp (D) or Arg (K). Helix breaking as Pro (P) or Gly (G). Helix forming: Leu (L), Ile (I), Phe (F), Glu (E), Tyr (Y), Trp (W), Met (M). Aromatic: Phe (F), Tyr (Y), Trp (W), His (H). Residue proportions within exons and in the flanking regions were defined by counts. The flanking regions were defined as residues 30 amino acids upstream and 30 amino acids downstream of the inserted exon. Enrichment was calculated as 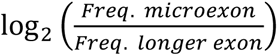, and statistically examined using fisher-exact test. Residue analysis of microexon overall as well as microexon subsets (microexons in helices, microexons in secondary structure) was compared to longer neural cassette overall.

#### Protein structural modelling

SNX2 isoforms from human, rat, chicken, and zebrafish were modelled using AlphaFold2 v2.1 with the default input settings through the application that is set up via Google Colaboratory^87^. The isoform models were visualised in ChimeraX^88^ v1.8rc202405240417. ChimeraX was also used to align, examine surface charge, and examine electrostatic interactions (only salt bridges) of the SNX2 region corresponding to 419 – 464 of human SNX2 (Uniprot ID: O60749) with default input settings.

Sequences for TEAD1 and ERC1 protein isoforms were extracted from UniProt database^89^ (human TEAD, Uniprot ID: P28347-1. Human ERC1, Uniprot ID: Q8IUD2-1) and amino acid sequences corresponding to the microexons-of-interest were added/removed manually. Protein structures were modelled using AlphaFold3 on the AlphaFold server^40^ (https://alphafoldserver.com) with default settings. Models were downloaded and visualized using ChimeraX.

#### Amino acid conservation analysis of the SNX2 microexon

All isoforms of SNX2 annotated on the UniProt server^89^ were identified and isoforms containing the microexon and of canonical size (∼500 amino acids) were extracted. Isoforms were further manually filtered to ensure no evolutionary clade was over-represented. Sequences were aligned using Clustal Omega (v1.2.4)^90^ using default parameters. JalView was then used to visualize sequences^91^. Secondary structure was predicted using JPred secondary structure prediction (Lupas_14) and JNetPRED, buried residues were predicted using Jnet burial prediction of solvent accessibility, and conservation using JNETConf^91^.

#### Interface-resolved interactome

Protein–protein interactions (PPIs) were mapped to domain–domain and motif–domain contacts based on curated interface annotations from Interactome3D^92^, Domino^93^, IntAct^94^, 3DID^95^ and ELM^96^. Only interactions with experimentally verified interface mappings were retained, producing an *interface-resolved interactome* in which each edge corresponds to a specific physical contact rather than a generic protein–protein association.

#### Identification of alternative splicing events

Vast-tools compare was used to analyse alternative splicing from 75 tissue and cell types (https://vastdb.crg.eu/wiki/Downloads)^24^ using default setting except --min_dPSI 20 with all combinations of splicing events assessed. Only differential events annotated as exon skipping events (S, C1, C2, C3) whose length was divisible by 3 (i.e. frame-preserving) and identified as differential between multiple (>=2) tissues were considered in the interface-resolved interactome.

#### Integration of alternative splicing data

For each identified AS exon, Shannon entropy (H) and Gini index (G) were calculated from its percent spliced in (PSI) distribution across tissues:

Shannon Entropy:

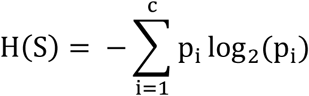

Gini Index:

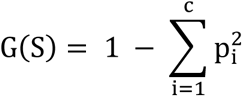

**-** G(S): Gini impurity of the set S
**-** H(S): Entropy of the set S
**-** c: The number of classes
**-** p_i_: The proportion (probability) of class *i* in the set

High-entropy and high-Gini exons correspond to tissue-specific splicing^97^, while low values reflect constitutive inclusion. For each in-frame exon its corresponding amino acids in the canonical ensembl protein transcript^98^ was identified. For the interface interactome, only those splicing events overlapping an annotated protein interface(s) were retained. Interactions regulated by these interfaces were considered AS-regulated edges. Alternative splicing events were annotated with additional information based on whether they overlapped known Pfam domain^99^, disordered region^100^, domain variety^99^ (RNA, DNA, protein, lipid), phosphorylation sites^101^ and uniport annotations^102^.

#### Percolation analysis

To evaluate the global impact of splicing on network stability, we progressively deleted interface-level edges in descending order *of AS regulation score* (i.e. removing the most tissue-specific exons first, then increasingly tissue-wide events). At each removal fraction f, we computed two measures:

- **ΔE** — the change in total number of protein–protein connections (degree sum).
- **ΔS** — the change in *network entropy*, reflecting redistribution of interactions across functional modules (computed as the Shannon entropy of community membership within connected components).

These values were compared to 1,000 degree-preserving random edge deletions to estimate empirical significance (p < 0.001). To assess whether observed network changes differed from random expectation, we implemented an empirical significance framework based on degree-preserving randomization. For each percolation experiment, 1,000 random replicates were generated in which edges were shuffled while maintaining each node’s degree distribution. At each step of progressive edge removal (fraction *f*), we recomputed ΔS (network entropy change) and ΔE (connectivity change) for both the observed and randomized networks. Empirical two-sided *p*-values were then calculated.

To summarize deviations across the entire percolation trajectory, we computed the area under each ΔS(*f*) or ΔE(*f*) curve (AUC) using the trapezoidal rule and compared observed AUCs to their null distributions using the same empirical approach. Reported *p*-values therefore reflect the proportion of randomizations exceeding the magnitude of the observed deviation in ΔS or ΔE, providing a non-parametric estimate of statistical significance.

The same edge removal sequence was applied in the inter-community edge analysis, in which we quantified the number of edges connecting distinct network modules (as defined by Louvain clustering). At each percolation step, the fraction of edges connecting nodes in different communities was calculated. This “inter-community edge fraction” (intercomm) quantifies the extent of cross-talk between modules. Changes in intercomm during edge removal were compared to 1,000 degree-preserving null networks, yielding empirical *p*-values for each *f* and for the AUC of the intercomm(*f*) curve. This revealed whether tissue-specific splicing preferentially affects inter-modular versus intra-modular links.

#### Ordinary Least Squares (OLS) Regression Analysis

To evaluate the determinants of alternative splicing (AS)–driven network variation, we fitted multiple ordinary least squares (OLS) regression models using standardized predictors derived from the interface-resolved interactome. All continuous variables were z-scored prior to modelling to ensure comparability of effect sizes and to minimize issues of scale. Model parameters were estimated using the *statsmodels* Python package (v0.14), and heteroscedasticity-robust (HC3) standard errors were applied in all cases.

Each model examined how structural and regulatory features—such as PFAM modularity, domain variety, intrinsic disorder, regulation score, and network centrality measures (e.g. betweenness, shared neighbours, clustering coefficients)—influence network outcomes including ΔS (network entropy change) and ΔE (edge loss). The dependent variable was fitted as a continuous outcome, with model fit evaluated using adjusted *R²* and model significance assessed by robust *F*-tests.

To test whether the effects of these predictors differed for microexon-containing proteins, we included explicit interaction terms (e.g. *PFAM×micro* or *regulation×micro*). For each interaction, both variables were mean-centered prior to multiplication to reduce collinearity. The resulting interaction term quantifies whether the slope of a continuous predictor (such as domain modularity or tissue-specific regulation) differs between microexon-containing and non-microexon proteins. In practical terms, this approach identifies whether microexons modulate the relationship between structural complexity and network connectivity. For binary variables such as *microexon status* (1 = microexon-containing, 0 = non-microexon), a significant interaction indicates that the predictor’s effect on the response variable (e.g. ΔS or inter-community connectivity) changes specifically in the presence of microexons. When significant interactions were detected, simple-slope analyses were conducted to estimate the direction and magnitude of the relationship in each group independently. Model coefficients and 95% confidence intervals were exported in tab-delimited format for visualization in coefficient (forest) plots. These plots display both the effect size and significance of each predictor, with interaction terms providing insight into how microexons reshape the relationship between structural features and network rewiring.

#### Localisation Entropy Analysis

To examine spatial consequences of these network effects, subcellular localisation data were obtained from the Human Protein Atlas^27^, where each protein was annotated with 1–4+ cellular compartments. A localisation entropy metric was computed for each protein as the Shannon diversity of its annotated compartments. Higher localisation entropy indicates a broader or more dispersed subcellular distribution. OLS models were then used with localisation entropy as the dependent variable and intercomm, microexon status, and their interaction (intercomm×micro) as predictors. This allowed testing whether microexons occurring at inter-community connectors modulate spatial dispersion. A positive coefficient indicates broader localisation, whereas a significant negative term indicates event restricts localisation to specific compartments.

#### Protein Annotation

Multi-module proteins are defined as proteins with multiple unique Pfam domains^99^. Phosphorylation sites were based on phosphosite annotation^101^. Proteins with interaction-substrate diversity are based on following Pfam annotation^99^ of domain types: **protein** (PF00017, PF00018, PF00397, PF00595, PF08416, PF01030, PF00498, PF00533, PF00244, PF00568, PF02213, PF00169, PF00373, PF01344, PF00400, PF00515, PF00514, PF02985, PF00023, PF00560, PF00651, PF00531, PF01335, PF00619, PF02758, PF00452, PF02809, PF00627, PF00789, PF00567, PF00385, PF00439, PF02820, PF01426, PF00622, PF00630), **DNA** (PF00096, PF00105, PF00046, PF00157, PF00172, PF00010, PF00170, PF00505, PF00250, PF00178, PF02319, PF08779, PF00249, PF00251, PF00907, PF00319, PF01388, PF00651, PF00554, PF04299, PF00126, PF00440, PF02037), **RNA** (PF00076, PF00013, PF00035, PF00270, PF00271, PF00806, PF02171, PF02170, PF01423, PF01479, PF01299, PF04146, PF00642, PF00098, PF00575, PF01336, PF02926, PF01472, PF01918, PF04352, PF01535), **lipid-binding** (PF00169, PF00168, PF01363, PF00787, PF00130, PF01417, PF07653, PF03114, PF10455, PF00611, PF00373, PF02149, PF02893, PF01167, PF01852, PF01237, PF00650, PF08718, PF00191, PF01477, PF00400). Gene annotations for Figure 1H were based on GO-term annotations downloaded from Ensembl Biomart database^98^.

### Statistics

Data was analysed and statistically tested as indicated in the sections relevant to said data. When multiple tests were performed, p-values were corrected for false discovery rate using the Benjamini–Hochberg Method. The exact number of experimental repeats are stated in the Figure Legends and/or Methods relating to the specific experiment. P-values ≤0.05 were considered significant for LUMIER and correlation tests, while P-values ≤0.01 were considered significant for the examination of normalised classifier performances. For network analysis, randomization and regression models used fixed random seeds for reproducibility with all reported *p*-values being two-sided unless otherwise specified, and empirical *p*-values are derived from 1,000 randomizations.

### Reagents, plasmids, and oligonucleotides

**Table 3.** List of used reagents.

| Reagents | Provider | Catalogue # |
| --- | --- | --- |
| **MOLECULAR CLONING** |  |  |
| Gateway™ LR Clonase™ II Enzyme mix | ThermoFisher | 11791020 |
| Gateway™ BP Clonase™ II Enzyme mix | ThermoFisher | 11789020 |
| Q5® Site-Directed Mutagenesis Kit | New England Biolabs | E0554S |
| NEB® 5-alpha Competent E. coli (High Efficiency) | New England Biolabs | C2987H |
| One Shot™ Stbl3™ Chemically Competent E. coli | ThermoFisher | C737303 |
| Wizard® Plus SV Minipreps DNA Purification Systems | Promega | A1330 |
| NucleoBond Xtra Midi kit for transfection-grade plasmid DNA | Macherey-Nagel | 740410 |
| PureYield™ Plasmid Maxiprep System | Promega | A2392 |
| **RT-PCR** |  |  |
| TRIzol reagent | ThermoFisher | 15596026 |
| Chloroform | SigmaAldrich | C2432 |
| QIAGEN OneStep RT-PCR kit | QIAGEN | 210212 |
| GelRed® Nucleic Acid Gel Stain | Biotium | #41003 |
| **MAMMALIAN CELL CULTURE** |  |  |
| DMEM, high glucose, pyruvate | ThermoFisher | 11995065 |
| Fetal Bovine Serum (FBS) | ThermoFisher |  |
| Penicillin-Streptomycin (10,000 U/mL) | ThermoFisher | 15140122 |
| Trypsin-EDTA (0.25%), phenol red | ThermoFisher | 25200056 |
| Blasticidin S HCl (10 mg/mL) | ThermoFisher | A1113903 |
| Lipofectamine™ 3000 Transfection Reagent | ThermoFisher | L3000001 |
| ViaFect™ Transfection Reagent | Promega | E4981 |
| **IMMUNOFLUORESCENCE** |  |  |
| Precision cover glasses thickness No. 1.5H (tol. $\pm 5 \mu\text{m}$) | Marienfeld | 0117580 |
| 24 Well glass bottom plate with high performance #1.5 cover glass | Cellvis | P24-1.5H-N |
| DAPI (4',6-diamidino-2-phenylindole, dihydrochloride) | ThermoFisher | D1306 |
| Golgin-97 Monoclonal Antibody (CDF4), eBioscience™ | ThermoFisher | 14-9767-82 |
| Alexa Fluor® 488 AffiniPure™ Fab Fragment Goat Anti-Mouse IgG (H+L) | Jackson ImmunoResearch | 115-547-003 |
| Alexa Fluor™ 647 NHS Ester (Succinimidyl Ester) | ThermoFisher | A20006 |
| Phalloidin-iFluor 647 Reagent | Abcam | ab176759 |
| Fluoromount-G™ Mounting Medium | ThermoFisher | 00-4958-02 |
| **CELL LINE GENERATION AND IMMUNOPRECIPITATION** |  |  |
| BD Transduction Laboratories™ Purified Mouse Anti-SNX2 | BD Biosciences | 611308 |
| GAPDH Antibody (0411): sc-47724 | Santa Cruz Biotechnology | sc-47724 |
| Goat anti Mouse IgG (H+L) Secondary Antibody, HRP | ThermoFisher | 31430 |
| DTT (dithiothreitol) | ThermoFisher | R0861 |
| NuPAGE™ LDS Sample Buffer (4X) | ThermoFisher | NP0007 |
| Bolt™ Bis-Tris Plus Mini Protein Gels, 8%, 1.0 mm, WedgeWell™ format | ThermoFisher | NW00085BOX |
| Immobilon®-P PVDF Membrane | SigmaAldrich | IPVH00010 |
| Clarity Western ECL Substrate | BioRad | #1705061 |
| Dynabeads™ Protein A for Immunoprecipitation | ThermoFisher | 10001D |
| Dynabeads™ Protein G for Immunoprecipitation | ThermoFisher | 10003D |
| Halt™ Protease and Phosphatase Inhibitor Single-Use Cocktail (100X) | ThermoFisher | 78442 |
| **LUMIER** |  |  |
| Monoclonal ANTI-FLAG® M2 antibody produced in mouse | SigmaAldrich | F1804-50UG |
| Monoclonal ANTI-FLAG® M2-Peroxidase (HRP) antibody produced in mouse | SigmaAldrich | A8592 |
| Renilla-Glo® Luciferase Assay System | Promega | E2710 |
| SuperSignal™ ELISA Pico Chemiluminescent Substrate | ThermoFisher | 37070 |
| Microplate, 96 Well, Ps, F-Bottom, (Chimney Well), White, Lumitrac, High Binding, Sterile | Greiner | 655074 |

**Table 4.** Used Addgene plasmids.

| Plasmid | Addgene catalogue # | Reference |
| --- | --- | --- |
| pSpCas9(BB)-2A-GFP (PX458) | Addgene (Plasmid #48138) | [PMID: 24157548] |
| pCAG-VSVG | Addgene (Plasmid #35616) | Unpublished |
| psPAX2 | Addgene (Plasmid #12260) | Unpublished |
| pLX303 | Addgene (Plasmid #25897) | [PMID:21706014] |
| pLenti CMV Puro DEST (w118-1) IGF1R | Addgene (Plasmid #107499) | PMID: 30404791 |
| R777-E106 Hs.INSR-nostop | Addgene (Plasmid #70390) | Unpublished |
| pcDNA3-hL1 | Addgene (Plasmid #89411) | [PMID: 10469653] |
| pDONR223_CXCR4_WT | Addgene (Plasmid #81957) | [PMID: 27147599] |
| pDONR221_SLC6A9 | Addgene (Plasmid #132220) | [PMID: 32265506] |
| pDONR221_SLC9A6 | Addgene (Plasmid #132175) | PMID: 32265506] |

**Table 5.**
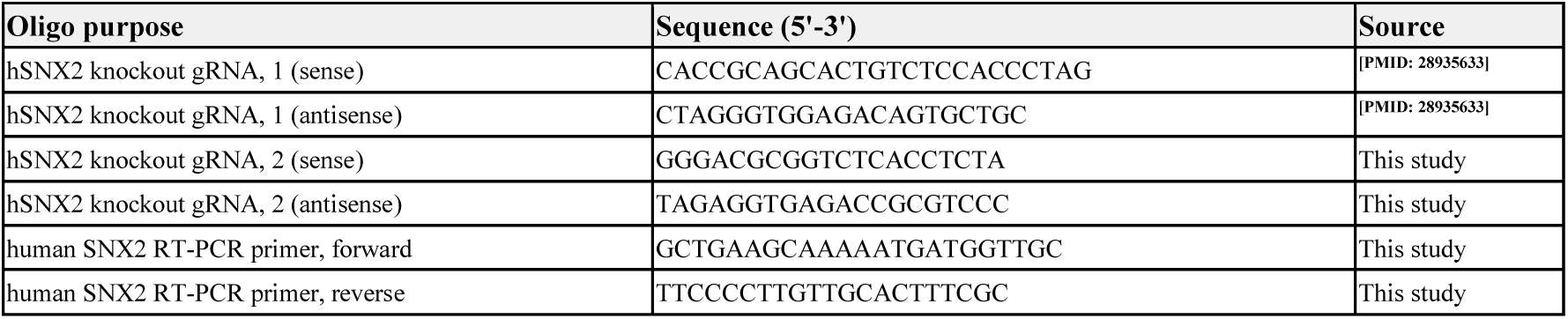
Key oligonucleotides.

## DATA AVAILABILITY

The imaging-based isoform library cell measurements have been deposited at Figshare: https://doi.org/10.6084/m9.figshare.29486819

The performance metrics of the classifiers generated from analysing the isoform cell measurements have been deposited at Figshare: https://doi.org/10.6084/m9.figshare.29487851

The classifier-derived feature importances for the isoform classifiers have been deposited at FigShare: https://doi.org/10.6084/m9.figshare.29487845

The imaging-based SNX2 mutant library cell measurements has been deposited at Figshare: https://doi.org/10.6084/m9.figshare.29487671

## CODE AVAILABILITY

In-house developed custom analysis scripts are available at https://github.com/weatheritt2/Hansen_AS/ under MIT license.

---

**Figure S1.**
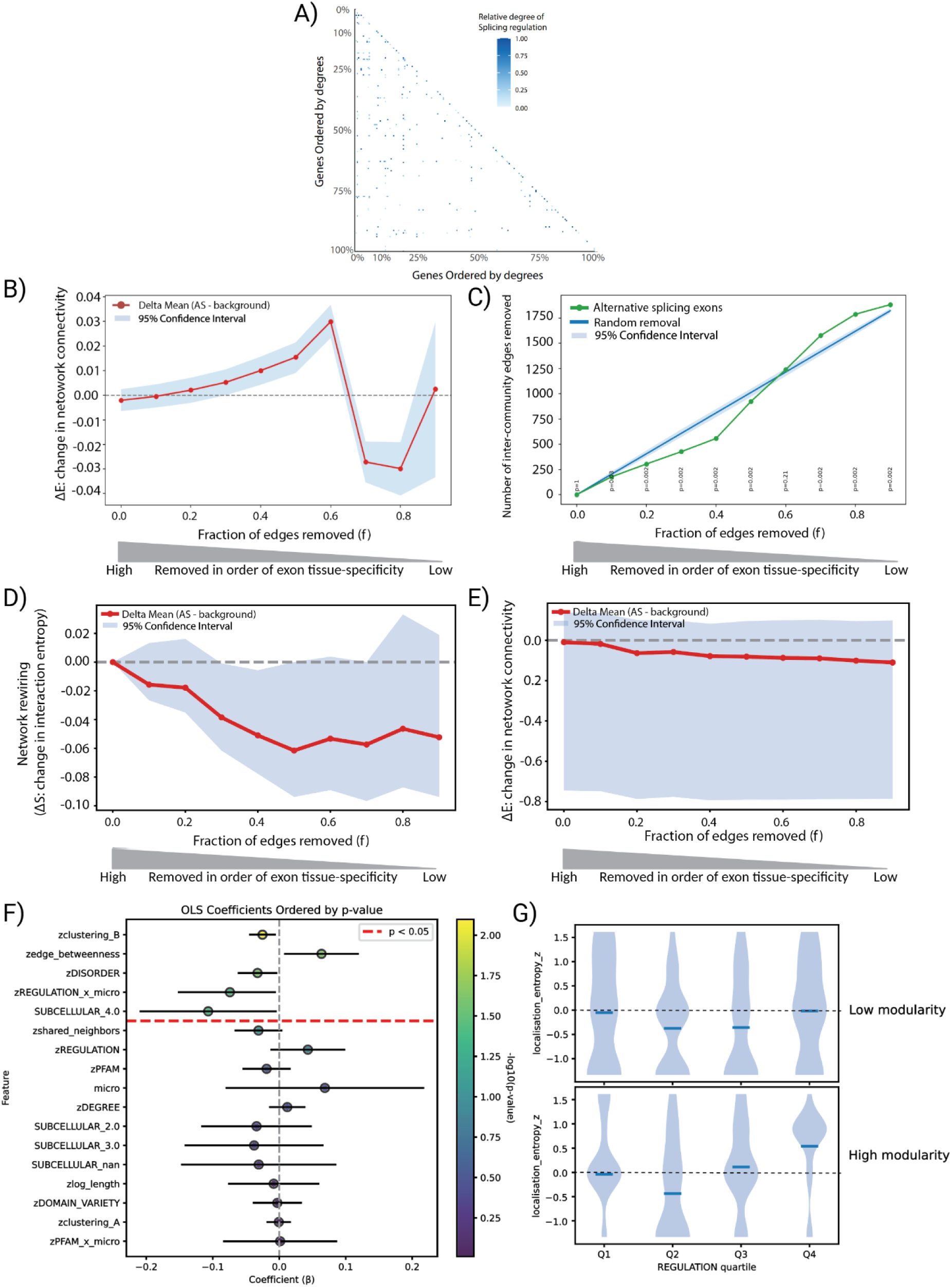
Alternative splicing-regulated interactions in protein network analysis. **A)** Triangular adjacency heatmap of the interface-resolved interactome ordered by total node degree (top 200 proteins with most interactions). Cells in heatmap denote protein-protein interfaces; colour encodes interaction diversity (network entropy; percentile scaled), with non-AS edges in white. **B)** Percolation analysis of network connectivity (ΔE) during progressive removal of AS-regulated edges. The red line indicates the mean difference in network connectivity (Δ = EAS – Enull)) as a function of the fraction of AS-regulated edges (f) progressively removed, from tissue-specific to tissue-wide events, with shaded regions showing the 95% confidence interval. The grey dashed line denotes Δ = 0 (no difference). To define the background distribution, null edges correspond to non-AS interactions randomly selected to match the degree distribution of AS-regulated edges. Positive Δ values indicate higher network connectivity among AS-regulated interfaces than expected by chance. **C)** Percolation analysis of inter-community connectivity during progressive removal of AS-regulated interfaces. The green line shows the number of inter-community edges removed as a function of the fraction of AS-regulated edges (f) progressively removed from tissue-specific to tissue-wide events. The blue line denotes the degree-matched background null mean, with shaded areas indicating the 95% confidence interval derived from 1,000 permutations. Values above the null indicate that AS-regulated interfaces disproportionately connect distinct functional communities **D)** Percolation analysis of network entropy during progressive removal of microexon-regulated edges. The red line indicates the mean difference in network entropy (Δ = SAS – Snull) as a function of the fraction of AS-regulated edges (f) progressively removed, from tissue-specific to tissue-wide events, with shaded regions showing the 95% confidence interval. The grey dashed line denotes Δ = 0 (no difference). Null edges correspond to non-AS interactions randomly selected to match the degree distribution of AS-regulated edges. Positive Δ values indicate higher interaction diversity among AS-regulated interfaces. **E)** Percolation analysis of change in network connectivity (ΔE) during progressive removal of microexon-regulated edges. The red line indicates the mean difference in network connectivity (Δ = EAS – Enull) as a function of the fraction of AS-regulated edges (f) progressively removed, from tissue-specific to tissue-wide events, with shaded regions showing the 95% confidence interval. The grey dashed line denotes Δ = 0 (no difference). Null edges correspond to non-AS interactions randomly selected to match the degree distribution of AS-regulated edges. Positive Δ values indicate higher network connectivity among AS-regulated interfaces then expected by chance. **F)** Multivariate regression analysis identifying features that predict whether an interaction occurs between or within functional communities. Shown are standardized coefficients from an ordinary least-squares (OLS) model, ordered by statistical significance (–log₁₀ p-value). Each point is standardised regression coefficient with horizontal line representing 95% confidence interval. Positive coefficients indicate features associated with inter-community interactions, whereas negative coefficients correspond to features enriched within community modules. **G)** Violin plots showing the distribution of localisation entropy across quartiles of splicing regulation, stratified by protein modularity. Localisation entropy (y-axis, z-score) quantifies the number of distinct subcellular compartments in which each protein is detected (Human Protein Atlas).

**Figure S2.**
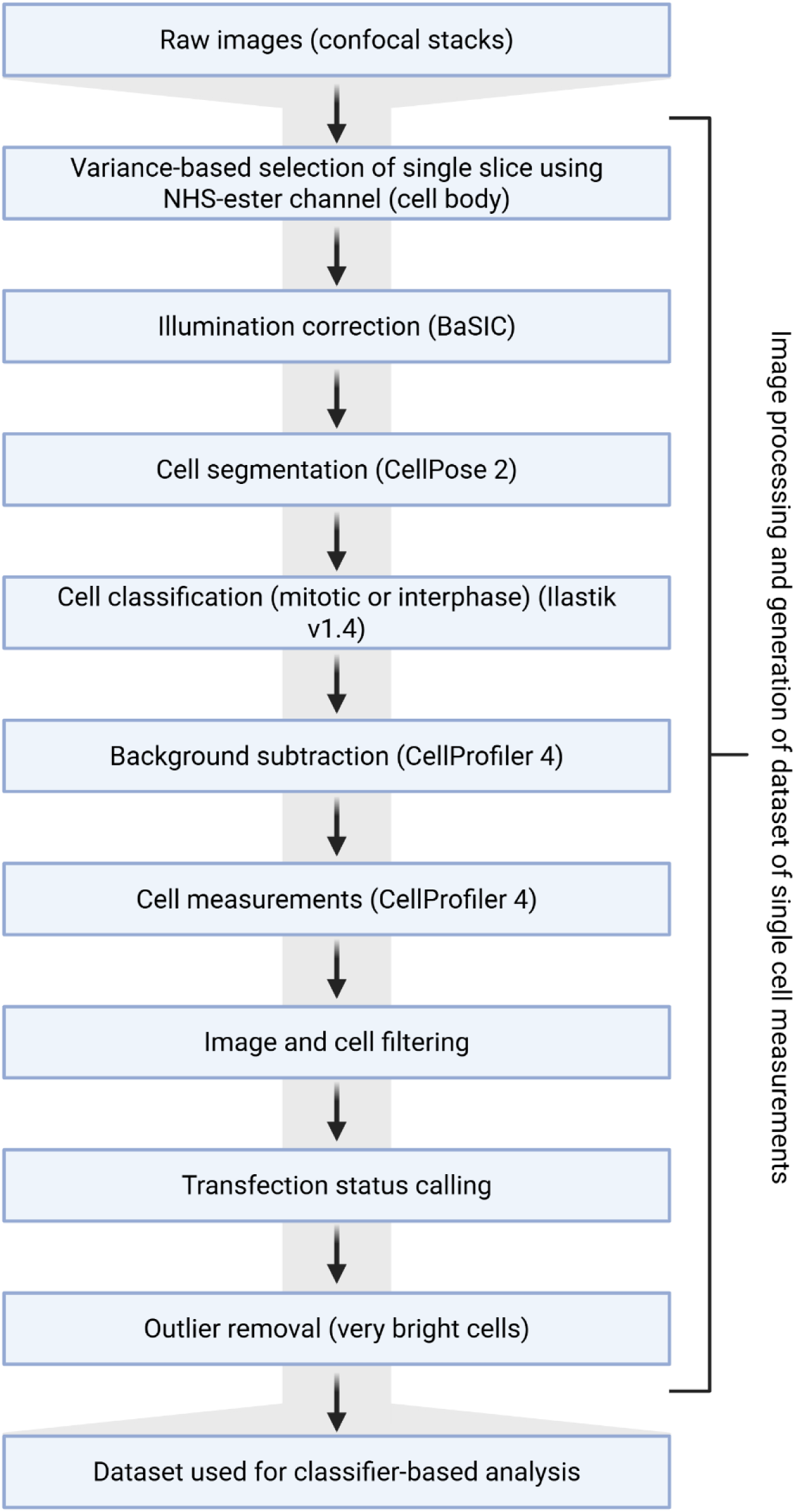
Generation of a single cell isoform measurement dataset from the acquired images. Overview of image processing to generate the cell datasets used for the classifier-based analysis (**Figure S3**). The selected focus slice from each confocal stack is used for all four channels for all subsequent processing. See Methods for details.

**Figure S3.**
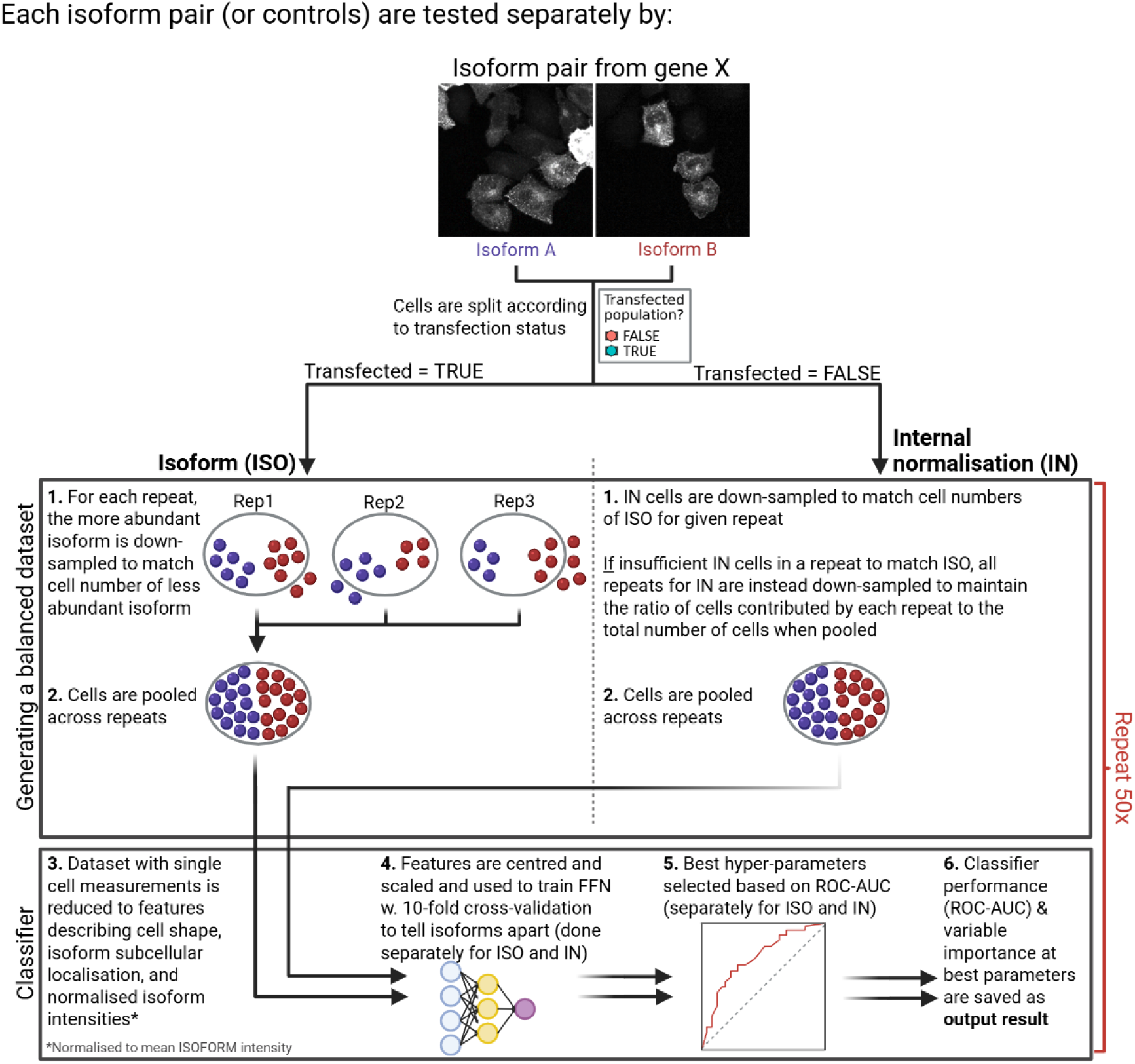
Isoform differences are detected using down-sampling and classifier-based analysis of single cell measurement dataset. Overview of classifier-based analysis. Isoforms (or the positive or negative control) are compared in a pairwise-manner, where isoform-expressing cells (transfected, referred to as ISO) and internal normalisation cells (non-transfected, referred to as IN) initially are split. For the ISO cells, the more abundant isoform is down-sampled to match the less abundant one in terms of cell numbers on a replicate basis (1). The cells are then pooled across replicates (2), which means replicates can contribute different number of cells, but always 50% of each isoform. This corrects for replicate-specific effects driving classification. The IN is sampled to match the ISO. The down-sampled datasets are then used for classifier-based analysis where features describing cell morphology, mCherry localisation, and normalised intensity values are used as input for a feedforward neural network (FFN) (3). The classifier is trained with 10-fold cross-validation where the average performance on the test folds at optimal parameters is used as a readout for classifier performance (4, 5). Feature importance scores are also saved to identify features driving classification (6). The classifier evaluation is done separately for the ISO and IN datasets. The combined down-sampling and classifier evaluation is done 50 times, where the performance for each iteration is used as the readout. See Methods for details.

**Figure S4.**
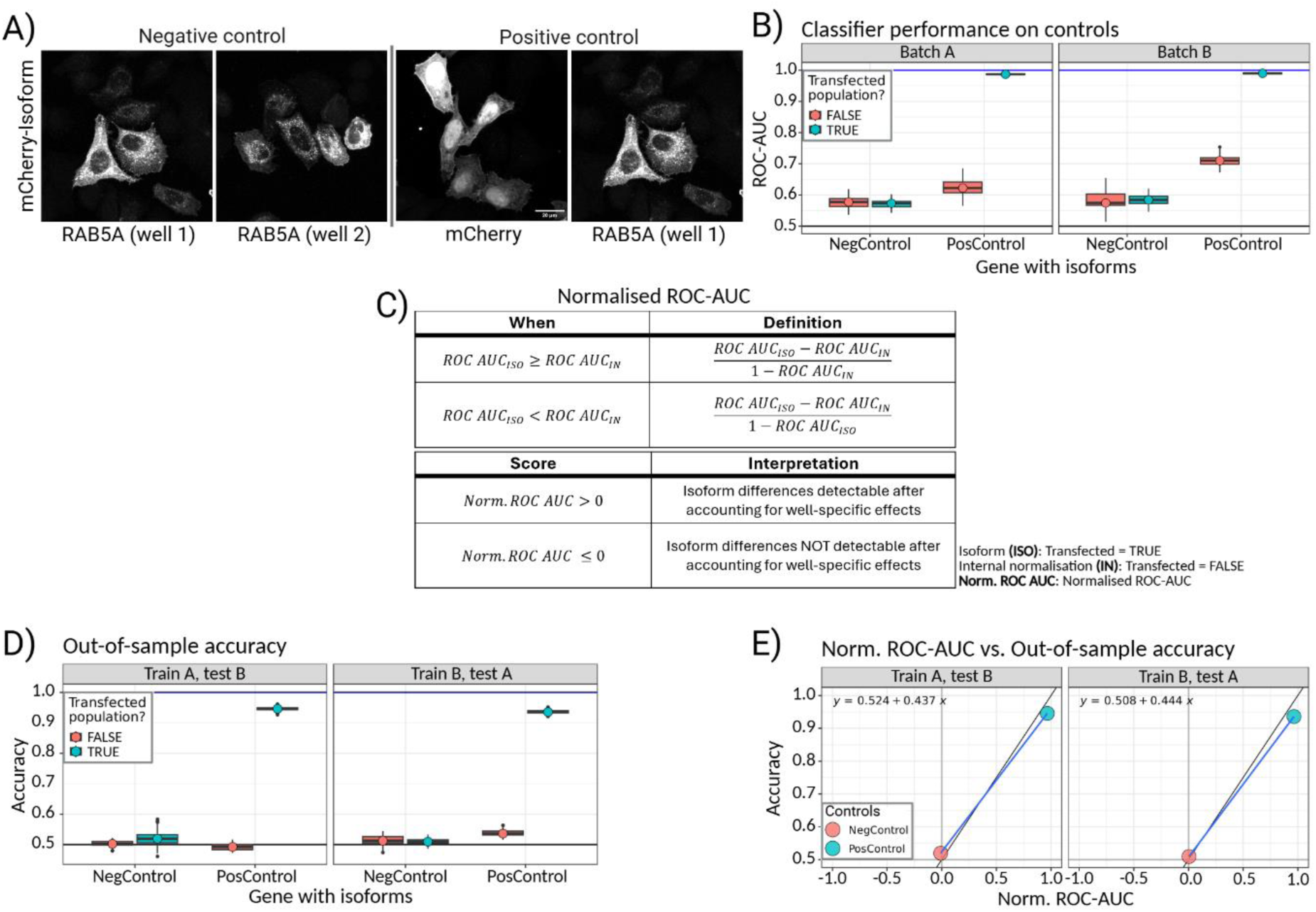
Validation of combined high-content imaging and classifier-based approach using well-defined controls. **A)** Example images of cells expressing the mCherry constructs used as positive and negative controls, same as **Figure 2D**. The negative control consists of two separate wells transfected with the same mCherry construct (mCherry-RAB5A). The positive control consists of two separate wells transfected with mCherry and mCherry-RAB5A, respectively. Scale bar is 20µm. **B)** Boxplot showing the ability of classifiers trained on mCherry image features from cells expressing control plasmids to correctly assign the plasmid ID (which corresponds to well ID), quantified as ROC-AUC (Receiver Operating Characteristics (ROC), Area Under the Curve (AUC)). Cells came from three experimental repeats that were pooled for the classifier-based analysis with n cells ≥ 500 for each comparison after pooling. The experiment was performed twice, batch A and batch B, that were evaluated separately. The boxplots illustrate the ROC-AUCs from each of 50 iterations of cell sampling and classifier evaluation, and dot indicates median ROC-AUC (n = 50). Classifier-based analysis was performed separately on transfected (ISO) and non-transfected (internal normalisation) cells from the same wells to assess contribution of well-specific effects on classification. Coloured lines indicate perfect (blue) and worst (black) possible performance, respectively. **C)** Definition and interpretation of the normalised ROC-AUC score, where the internal normalisation (IN) classifier (using non-transfected cells, Transfected = FALSE) is used to normalise the isoform classifier (consisting of transfected cells, Transfected = TRUE). **D)** Boxplot showing performance of classifiers trained on mCherry image features from cells transfected with control plasmids in batch A to predict plasmid identity in batch B and vice versa, as quantified by accuracy on the test batch (out-of-sample accuracy). The test batch (A or B) was down-sampled to have equal amounts of cells expressing either plasmid in a control pair before being used for prediction. The accuracy can therefore range from 0.5 to 1.0. 20 iterations of classifier training and testing was done, and the boxplots shows the mean accuracy from each iteration, and dot indicates median accuracy across iterations (n = 20). n cells ≥500 for each condition in both test and training sets. **E)** XY plot showing the correlation between the normalised ROC-AUC (norm. ROC-AUC) and the out-of-sample accuracy for the two controls. The normalised ROC-AUC for batch A is compared to the out-of-sample accuracy when trained on batch A and tested on B (vice versa for ROC-AUC for batch B). A linear regression was fitted to the data.

**Figure S5.**
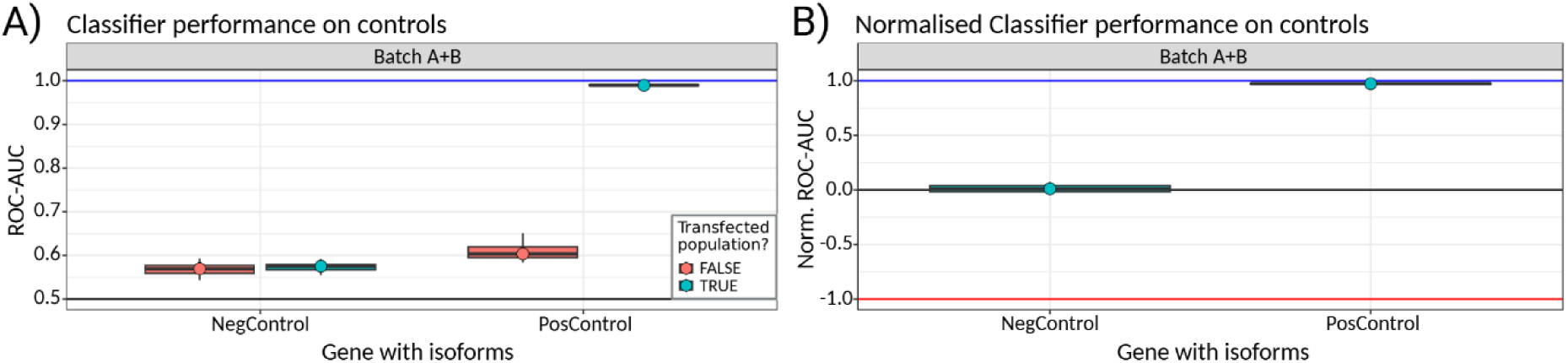
Pooling cells across experiment batches doesn’t alter performance for the positive and negative controls. **A)** Boxplots showing the ability of classifiers trained on mCherry features measured in cells expressing control plasmids to correctly assign the plasmid ID (which corresponds to well ID), quantified as ROC-AUC. Cells were pooled from two separate experiments (batch A and B), each with three experimental repeats, before the classifier-based evaluation (n cells ≥ 1500 after pooling). The boxplots illustrate the ROC-AUCs from each of 50 iterations of cell sampling and classifier evaluation, and dots indicate median ROC-AUC. Classifier-based analysis was performed separately on transfected (isoform) and non-transfected (internal normalisation) cells from the same, indicated by colour. Coloured lines indicate perfect (blue) and worst (black) possible performance, respectively. **B)** Same as **Figure S5A**, but the boxplots display classifier performance quantified as normalised ROC-AUC (norm. ROC-AUC). Coloured lines indicate perfect (blue), neutral (black) or worst (red) possible performance, respectively.

**Figure S6.**
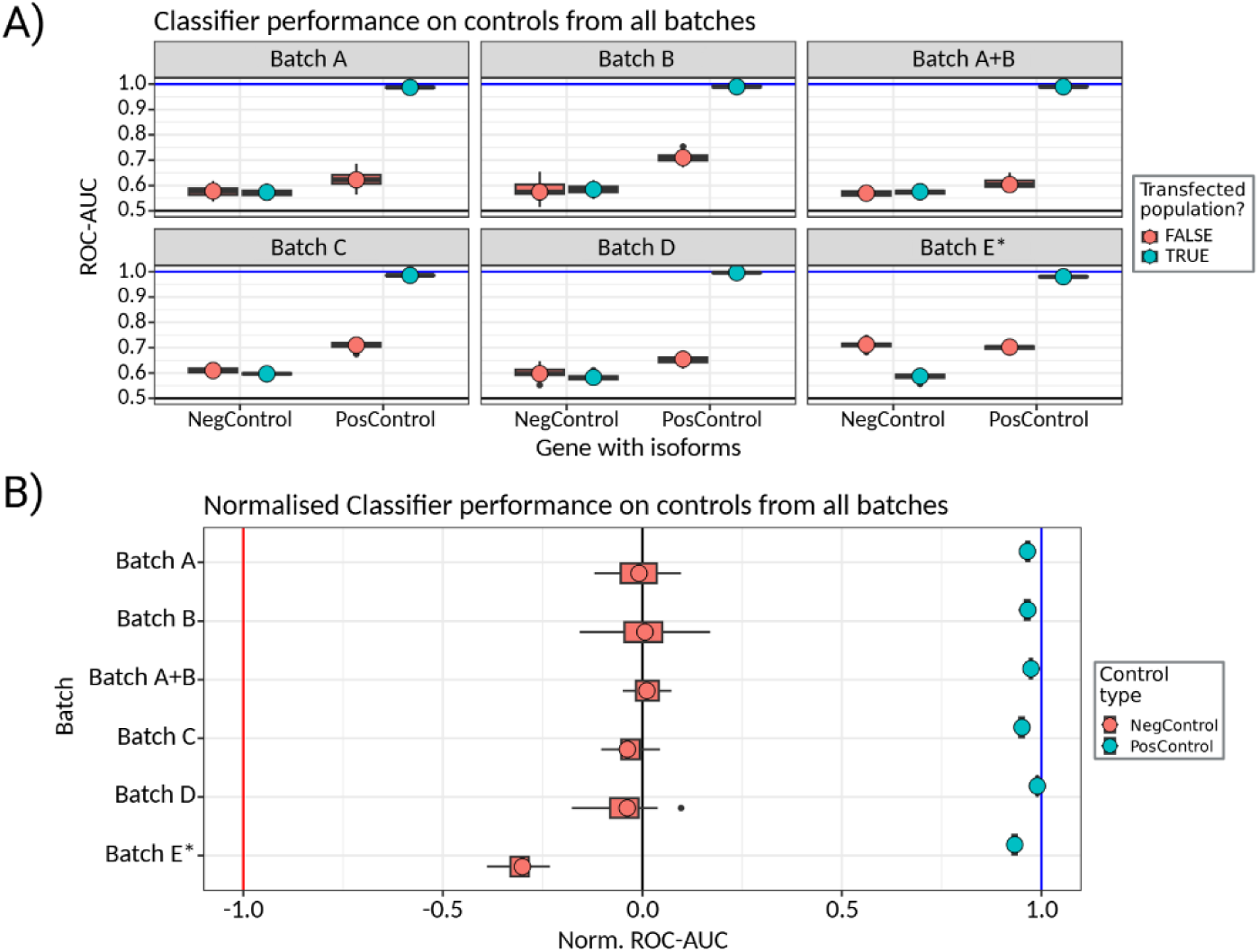
Performance on positive and negative controls is largely consistent across batches. **A)** Boxplots showing the ability of classifiers trained on mCherry features measured in cells expressing control plasmids to correctly assign the plasmid ID (which corresponds to well ID), quantified as ROC-AUC. The evaluation was performed separately on all experimental batches as well as the pooled batch A + B. Each batch consisted of three experimental repeats. n cells ≥ 500 for each comparison after pooling. The boxplots illustrate the ROC-AUCs from each of 50 iterations of cell sampling and classifier evaluation, and dots indicate median ROC-AUC across iterations. Classifier-based analysis was performed separately on transfected (isoform) and non-transfected (internal normalisation) cells from the same wells, indicated by colour. Coloured lines indicate perfect (blue) and worst (black) possible performance, respectively. * indicates Batch E. **B)** Same as **Figure S6A**, but the boxplots display classifier performance quantified as normalised ROC-AUC (norm. ROC-AUC). Coloured lines indicate perfect (blue), and neutral (black) performance, respectively.

**Figure S7.**
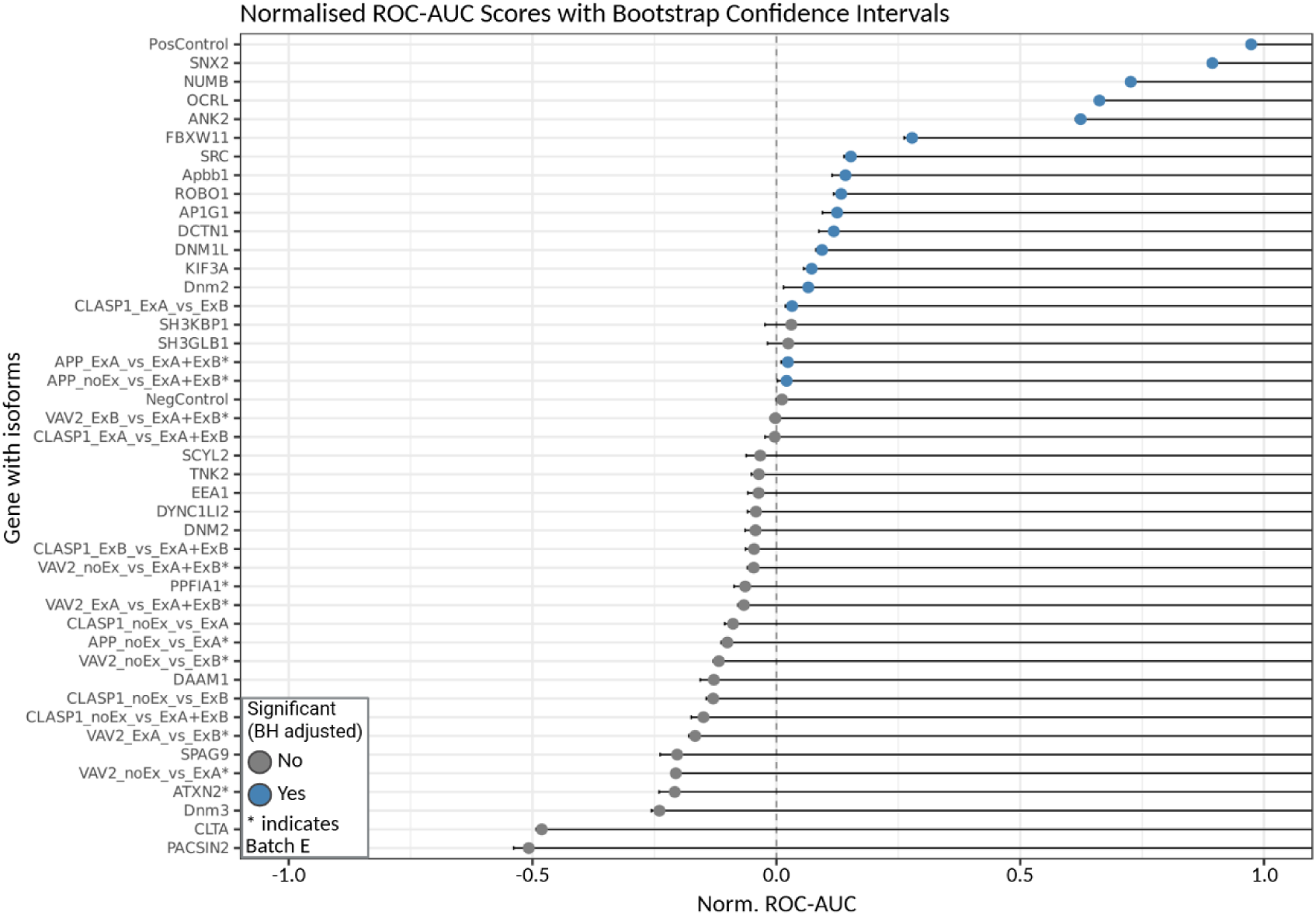
Bootstrap analysis of normalised ROC-AUCs. XY plots showing bootstrap statistical testing of the normalised ROC-AUC (norm. ROC-AUC) scores for the isoform pairs from **Figure 2F**. The normalised ROC-AUC scores for each pair were analysed via bootstrapping, and the displayed values are the bootstrap mean and 99% confidence intervals (CI). Isoform pairs that didn’t contain 0 in the lower bound of the CI after false discovery rate correction were identified as having ISO classifiers significantly outperforming the IN classifier, interpreted as isoform differences are detectable after considering well-specific effects, and are coloured blue. Isoform pairs with 0 in the CI were found as non-significant, coloured grey. n bootstrap iterations = 2000, one-sided test, p < 0.01, *.

**Figure S8.**
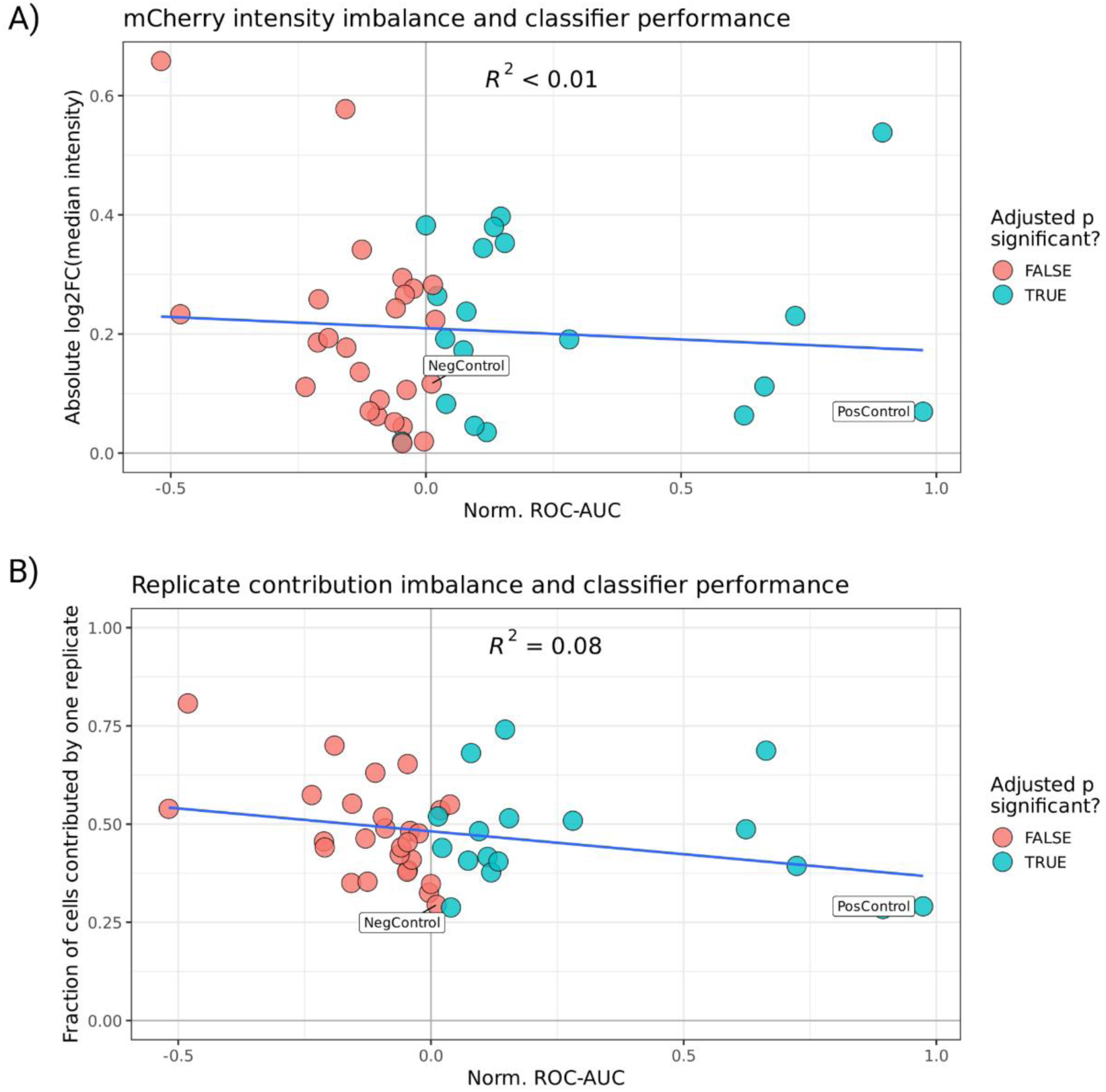
Classifier performance is not driven by population-level differences in expression amounts of isoforms within pairs nor by imbalanced contributions of cells form single replicates. **A)** XY plot of normalised classifier performance (norm. ROC-AUC) against expression imbalance between isoforms within the isoform pair (calculated as absolute log_2_(median intensity_+EXON/median_intensity_-EXON) for all cells used for classifier assessment). Each dot represents an isoform pair (n = 42). Dot colour indicates whether isoforms were found to be differentially localising or not. Positive and negative controls are labelled. A linear regression line (x ∼ y) is fitted to the data to describe the correlation. **B)** XY plot of normalised classifier performance (norm. ROC-AUC) against how large a fraction of cells originated from one replicate. Each dot represents an isoform pair (n = 42). Dot colour indicates whether isoforms were found to be differentially localising or not. Positive and negative controls are labelled. A linear regression line (x ∼ y) is fitted to the data to describe the correlation.

**Figure S9.**
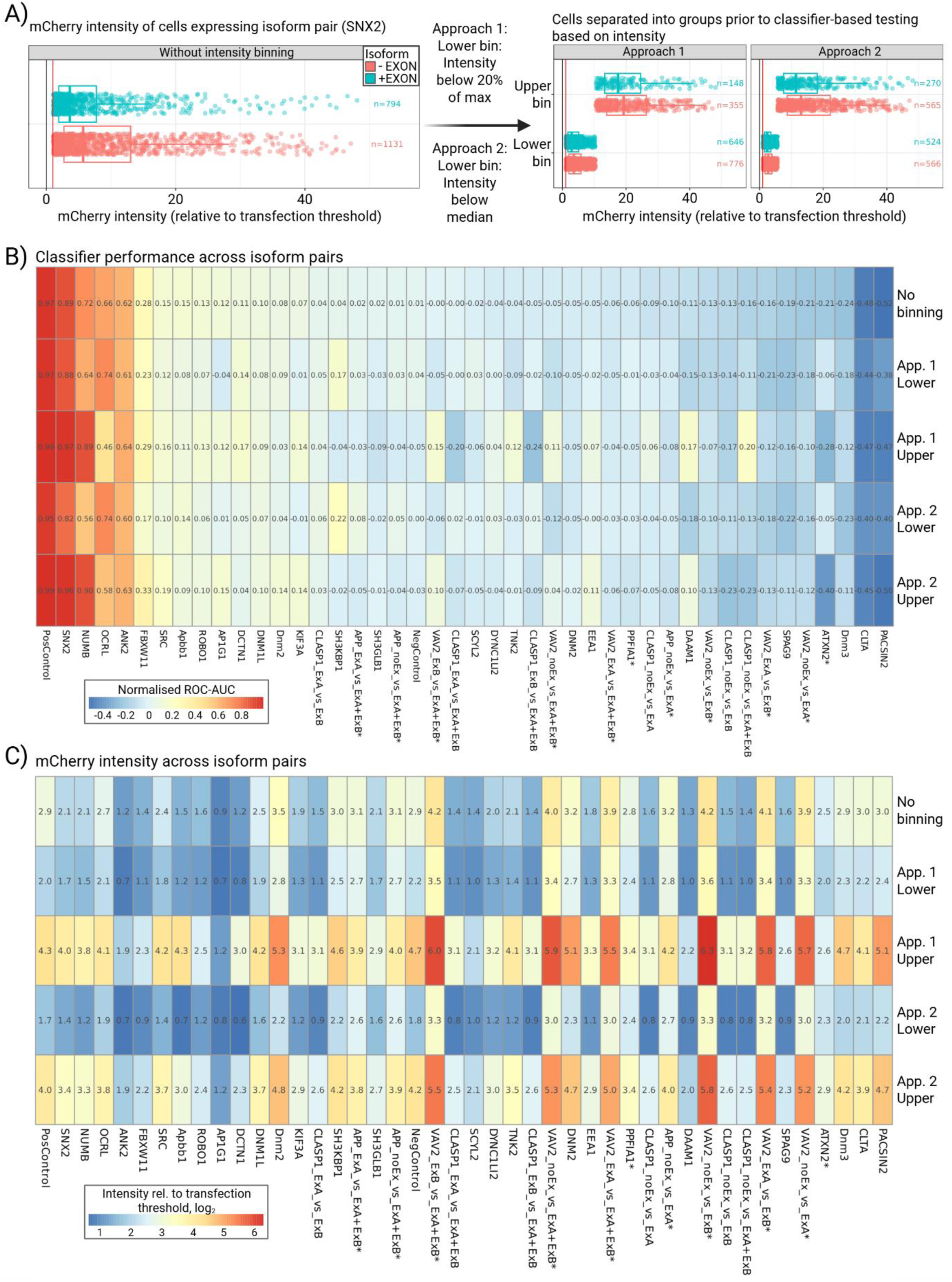
Separating cells in isoform pairs into high and low expression groups before classifier-based analysis yields comparable results to no separation. **A)** Boxplots of cell mCherry intensity (relative to the transfeciton threshold) of cells expressing mCherry-SNX2 isoforms with individual cells plotted as dots, before (**left**) and after (**right**) separating cells into low and high expression bins. Two binning approaches were used in parallel (Methods). Note, the binning thresholds were calculated for both isoforms, and the lower threshold was then applied across both isoforms. Colours indicate isoform identity. Red line indicates transfection threshold. **B)** Heatmap showing normalised classifier performance (norm. ROC-AUC) for all isoform pairs when using the full isoform dataset (No binning) or lower or higher intensity cells, separated via approach 1 or 2 outlined, as outlined in **Figure S9A**. App. = Approach. **C)** Heatmap showing median mCherry intensity relative to transfection threshold (log_2_ transformed), for cells expressing isoforms from a given gene, either using all cells (No binning) or after cells were separated into lower and higher intensity bins. App. = Approach.

**Figure S10.**
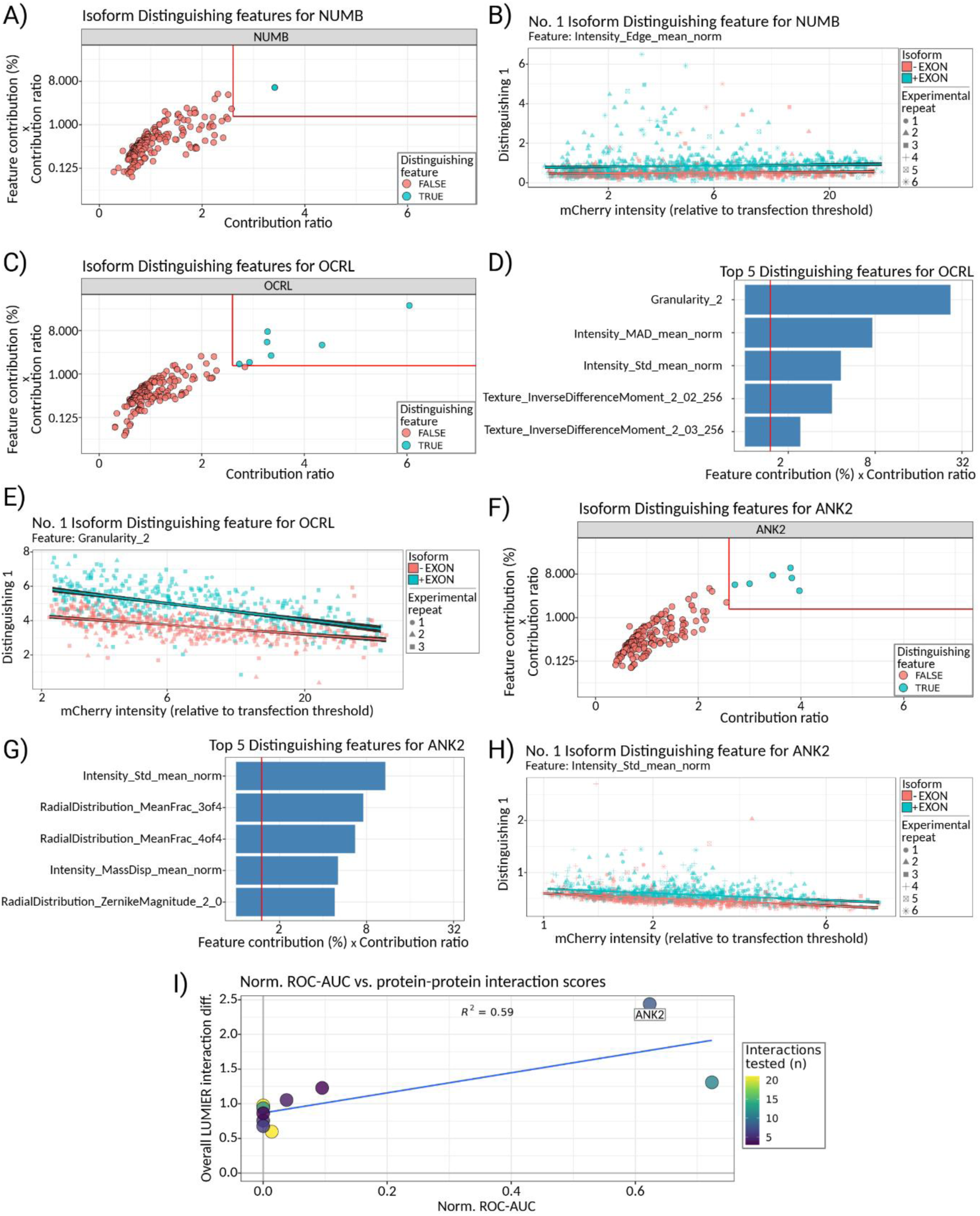
Isoform distinguishing features for NUMB, OCRL, and ANK2, and association between localisation and interaction. **A)** XY plot of Contribution ratio and Contribution ratio multiplied with Feature contribution (%) for the ISO classifier for each feature for the NUMB isoform pair, used to identify Isoform Distinguishing features (Methods). Of ∼200 features, 1 was identified as ‘Isoform distinguishing’. **B)** XY plot of Top 1 Isoform Distinguishing Feature for the NUMB isoforms (mean mCherry intensity at cell edge, normalised to mean mCherry intensity for whole cell) against cell mean mCherry intensity (relative to transfection threshold). Each data point corresponds to a cell with colour and shape indicating NUMB isoform identity and experimental repeat ID, respectively. Cells come from N = 6 experimental repeats, with pooled n cells > 1000. Separate linear regression lines were fitted to NUMB+EX and NUMB-EX-expressing cells. **C)** XY plot of Contribution ratio and Contribution ratio multiplied with Feature contribution (%) for the ISO classifier for each feature for the OCRL isoform pair, used to identify Isoform Distinguishing features (Methods). Of ∼200 features, 7 were identified as Isoform Distinguishing. **D)** Top 5 Isoform Distinguishing features according to their Feature contribution (%) multiplied with Contribution ratio for the OCRL isoform pair. The threshold value defining Isoform Distinguishing features is indicated as a red line. **E)** XY plot of Top 1 Isoform Distinguishing Feature for the OCRL isoforms (mCherry Granularity) against cell mean mCherry intensity (relative to transfection threshold). Each data point corresponds to a cell with colour and shape indicating OCRL isoform identity and experimental repeat ID, respectively. Cells come from N = 3 experimental repeats, with pooled n cells > 700. Separate linear regression lines were fitted to OCRL+EX and OCRL-EX-expressing cells. **F)** XY plot of Contribution ratio and Contribution ratio multiplied with Feature contribution (%) for the ISO classifier for each feature for the ANK2 isoform pair, used to identify Isoform Distinguishing features (Methods). Of ∼200 features, 6 were identified as Isoform Distinguishing. **G)** Barplot of top 5 Distinguishing features according to their Feature contribution (%) multiplied with Contribution ratio for the ANK2 isoform pair. The threshold value defining Isoform Distinguishing features is indicated as a red line. **H)** XY plot of Top 1 Isoform Distinguishing feature for the ANK2 isoforms (mCherry intensity standard deviation, normalised to mean mCherry intensity) against cell mean mCherry intensity (relative to transfection threshold). Each data point corresponds to a cell with colour and shape indicating ANK2 isoform identity and experimental repeat ID, respectively. Cells come from N = 6 experimental repeats, with pooled n cells > 900. Separate linear regression lines were fitted to ANK2+EX and ANK2-EX-expressing cells. **I)** XY plot examining the correlation between the normalised ROC-AUC (norm. ROC-AUC) and overall LUMIER interaction difference (a protein-protein interaction assay) (Methods) for isoform pairs where ≥3 interactors have been found to interact with at least one of the isoforms in a pair, n = 10 isoform pairs. A linear regression was fitted to the data, and the correlation was tested using a correlation test (cor(method = ’Kendall’)).

**Figure S11.**
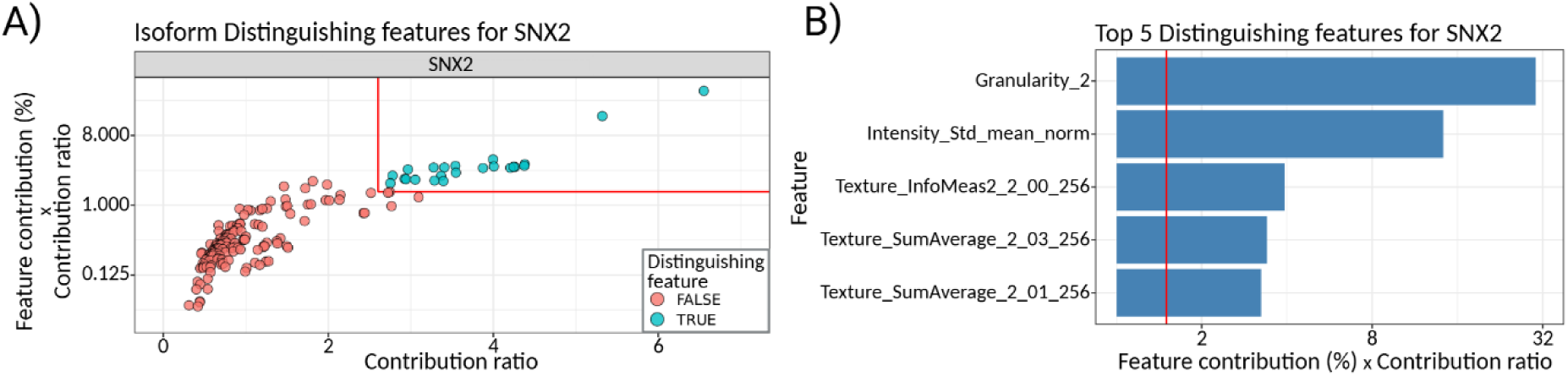
SNX2 isoform distinguishing features. **A)** XY plot of Contribution ratio and Contribution ratio multiplied with Feature contribution (%) for the ISO classifier for each feature for the SNX2 isoform pair, used to identify Isoform Distinguishing features (Methods). Of ∼200 features, ∼20 was identified as Isoform distinguishing. **B)** Barplot of the top 5 Isoform distinguishing features according to their Feature contribution (%) multiplied with Contribution ratio. The threshold value defining Isoform Distinguishing features is indicated as a red line.

**Figure S12.**
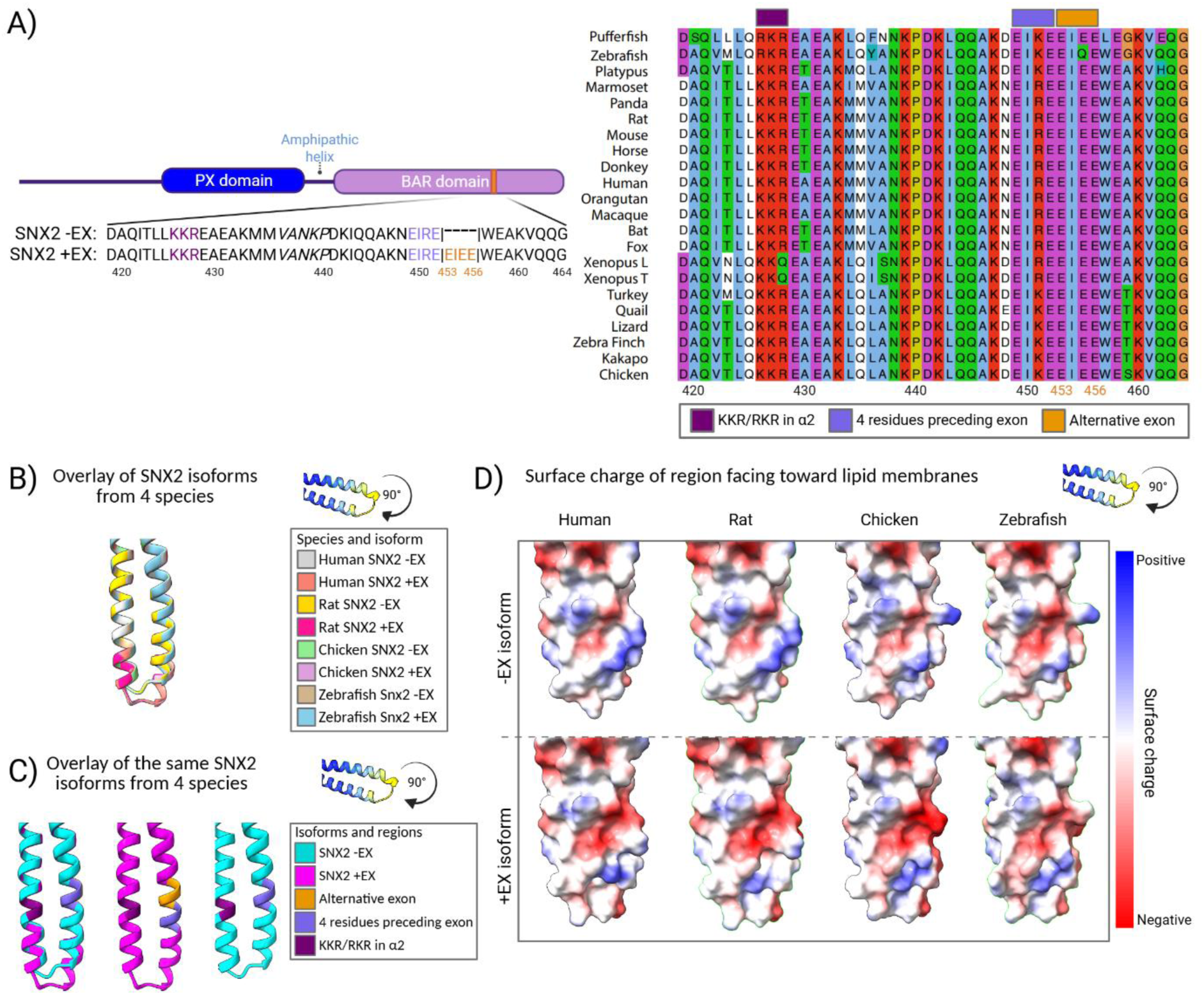
SNX2’s microexon is highly conserved, and is predicted to affect structure in a similar manner across vertebrates. **A)** Multiple sequence alignment of SNX2 region corresponding to human SNX2 residues 419-464 for 22 vertebrate species. The location of the microexon (orange), the 4 residues preceding the microexon (lavender), and the KKR motif (dark purple) are indicated by coloured rectangles. The individual positions are coloured according to the residue properties that is conserved across the aligned sequences (default clustal colouring from JalView). **B)** Overlay of AlphaFold2 models of the SNX2 BAR region containing residues 419-464 (human SNX2 residue numbering) of both SNX2 isoforms from four species (human, rat, chicken, zebrafish). The BAR domain is viewed from the side that faces lipid membranes. Each species-isoform combination has a distinct colour. **C)** Overlays of AlphaFold2 models of the SNX2 BAR region containing residues 419-464 (human SNX2 residue numbering) from four species (human, rat, chicken, zebrafish), separated according to isoform identity (±EXON). Colours reflect isoform (-EX cyan, +EX magenta), microexon (orange), the four residues preceding the microexon (lavender), and the KKR motif (dark purple), rather than species-isoform combination. **D)** Surface charge of AlphaFold2 models of SNX2 isoforms from the four species (human, rat, chicken, zebrafish), in the BAR region containing residues 419-464 (human SNX2 residue numbering). The BAR domain is viewed from the side that faces lipid membranes.

**Figure S13.**
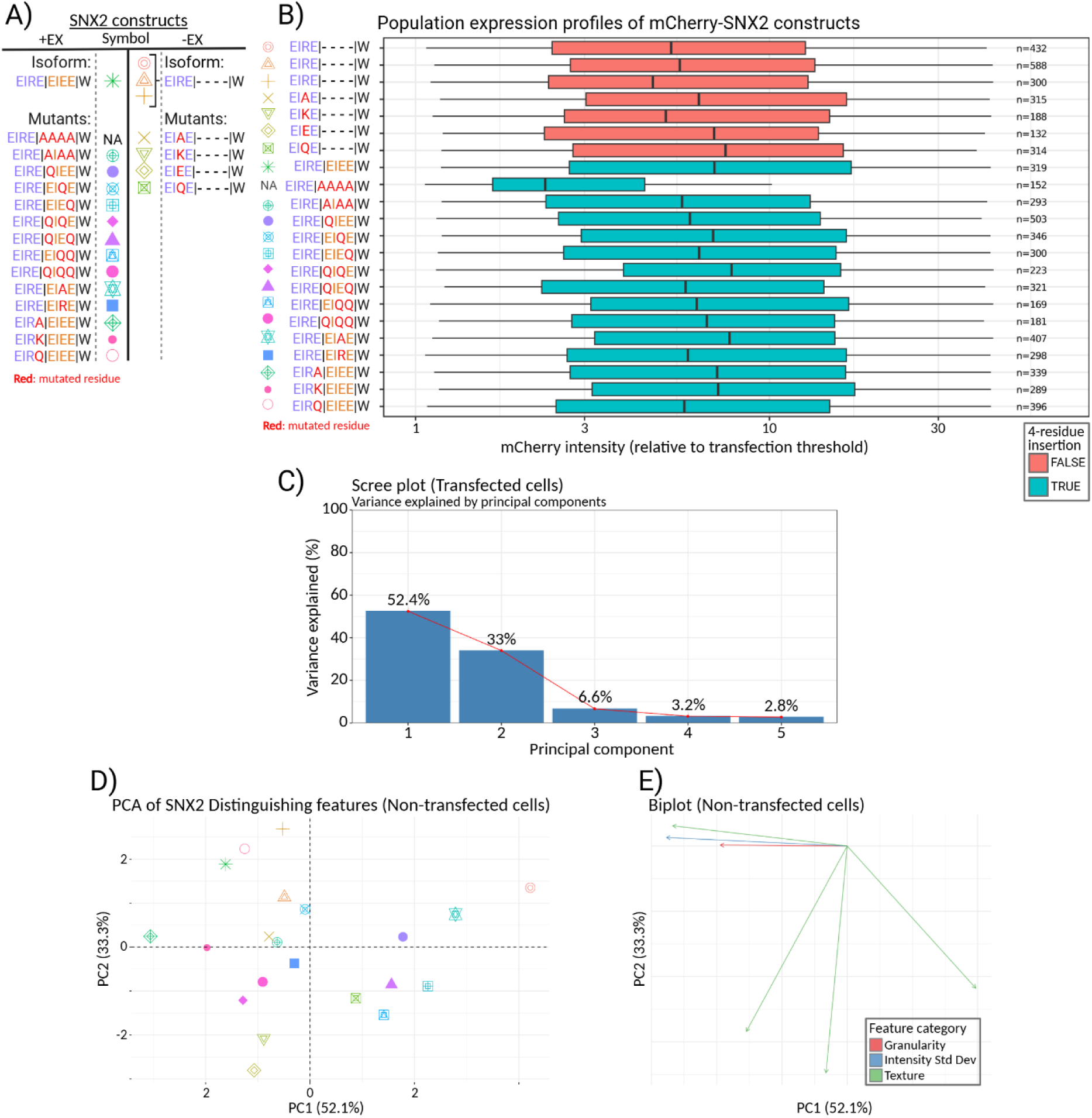
All but one mCherry-SNX2 constructs express at comparable levels and PCA clustering is driven by construct-dependent effects. **A)** List of SNX2 isoform and mutant constructs examined via high-content imaging. Same as **Figure 4F**. **B)** Boxplots showing cell mCherry mean intensity (relative to transfection threshold) of transfected cells expressing the various mCherry-SNX2 isoforms and mutants. Cells were pooled from three experimental repeats (N = 3). The number of cells after pooling is indicated for each construct (n). **C)** Scree plot showing principal component (PC) variance from PCA of SNX2 isoform-distinguishing features from transfected cells expressing SNX2 isoforms and mutants, relates to **Figure 4I**. **D)** XY plot showing PC1 versus PC2 from principal component analysis (PCA) of the mCherry-SNX2 construct imaging data using the population means of the SNX2 isoform-distinguishing features, measured for the non-transfected cells as input. **E)** Biplot showing the rotation of the Isoform-distinguishing features. Colours reflect feature class, with Granularity is shown in red.

**Figure S14.**
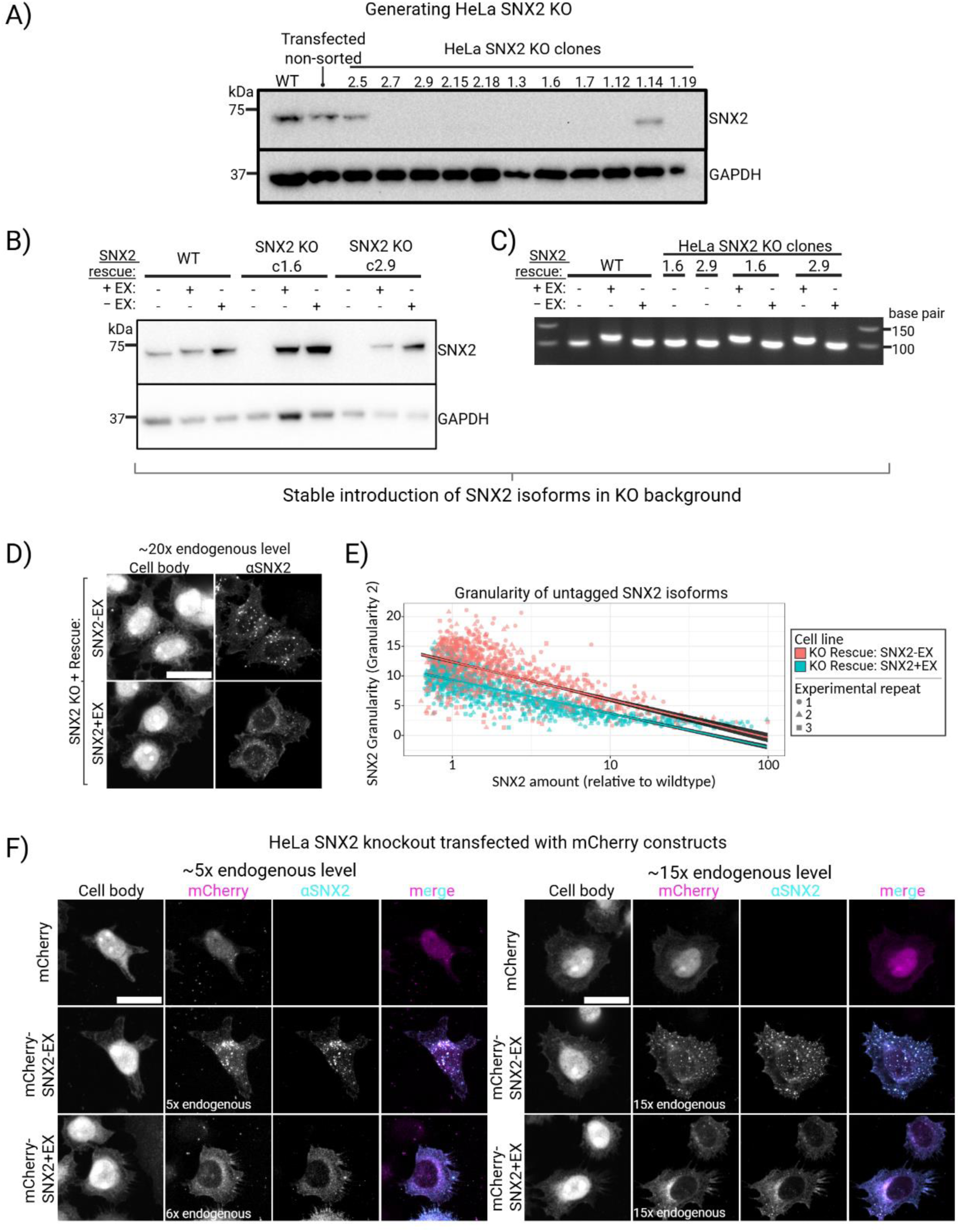
Generation of HeLa cell lines expressing select SNX2 isoforms. **A)** Western blot against human SNX2 and human GAPDH in lysates from HeLa cell clones isolated after SNX2 knockout in parental HeLa population. Experiment was performed once. **B)** Western blot against human SNX2 and human GAPDH in lysates from wild type HeLa cells or HeLa SNX2 knockout clones, where SNX2-EX or SNX2+EX had subsequently been re-introduced via lentiviral transduction. Experiment was performed once. **C)** End-point RT-PCR amplification of SNX2 region that flanks the SNX2 microexon in RNA from wild type HeLa cells or HeLa SNX2 lines. The PCR products were run on agarose gels, and a slight reduction in band mobility was visible upon microexon inclusion (12nt). The presence of PCR product in KO cells may stem from low amounts of SNX2 mRNA that contains the amplified region, as KO was directed at locus upstream of the amplified region. Experiment was performed once. **D)** Representative images of HeLa SNX2 rescue lines expressing SNX2 at ∼20x endogenous amounts with SNX2 detected via immunostaining. Scale bar, 25µm. **E)** XY plot of relative SNX2 amount (SNX2 intensity relative to median SNX2 intensity in HeLa wildtype) against SNX2 granularity 2 in the HeLa SNX2 rescue lines across a large expression range. Each point represents a single cell, colour represents cell line (n Rescue SNX2-EX = 1053, Rescue SNX2+EX = 960), and shape represents experimental repeat (N = 3). A linear regression line (x ∼ y) is fitted to the data, with the 95% confidence interval indicated as a black shade. **F)** Representative images of HeLa SNX2 KO cells transiently expressing mCherry constructs corresponding to ∼5x (**left**) or ∼15x (**right**) endogenous SNX2 amount with SNX2 detected via immunostaining. The actual expression level of the cells is indicated as white text. Scale bar: 25µm.

